# Conserved use of the sodium/bile acid cotransporter (NTCP) as an entry receptor by hepatitis B virus and domestic cat hepadnavirus

**DOI:** 10.1101/2023.02.04.527117

**Authors:** Maya Shofa, Akiho Ohkawa, Yasuyuki Kaneko, Akatsuki Saito

## Abstract

The *Orthohepadnavirus* genus includes hepatitis B virus (HBV) that can cause chronic hepatitis and hepatocarcinoma in humans. Recently, a novel hepadnavirus in cats, domestic cat hepadnavirus (DCH), was identified that is genetically close to HBV. DCH infection is associated with chronic hepatitis in cats, suggesting a similarity with HBV pathogenesis and the potential to use DCH as a novel animal model for HBV research. HBV is shown to use the sodium/bile acid cotransporter (NTCP) as a major cell entry receptor, but the equivalent receptor for DCH remains unknown. Here we sought to identify the entry receptor for DCH. HBV- and DCH-derived preS1 peptides efficiently bound to both human and cat NTCPs, and residue 158 of NTCP proteins determined the species-specific binding of the DCH preS1 peptide. Myrcludex B, an HBV entry inhibitor, blocked binding of the DCH preS1 peptide. Thus, DCH and HBV may share cell entry molecules, suggesting a possibility of inter-species transmission. Furthermore, our study suggests that DHC can be useful as a novel model for HBV research.

## 1. Introduction

*Hepadnaviridae* are enveloped DNA viruses that have a partially double-stranded, circular DNA genome and a broad host tropism, including mammals, reptiles, frogs, birds, and fish (Magnius et al., 2020). Human infection with hepatitis B virus (HBV), a member of *Orthohepadnavirus* genus, can induce chronic liver damage, including cirrhosis and hepatocellular carcinoma, and, despite effective vaccines being available, eradicating chronic infections can be challenging. Therefore, suitable HBV animal models are required to advance therapeutic outcomes. Chimpanzees are now unavailable as animal models for HBV infection, and infection models using HBV-like viruses in woodchuck, ground squirrel, and duck have consequently been investigated (Delmas et al., 2002; Guo et al., 2000; Jilbert et al., 1992; Kajino et al., 1994; Marion et al., 2002). Although treeshrews (Tupaia) are used as an animal model, the limited availability of research reagents may make it difficult to use this animal. Therefore, development of a novel and convenient animal model is still required.

In 2018, an HBV-like virus, domestic cat hepadnavirus (DCH), was identified in domestic cats in Australia and was associated with feline chronic hepatitis (Aghazadeh et al., 2018), suggesting a similarity with HBV pathogenesis. Subsequently, DCH was identified in other countries, including Italy (Lanave et al., 2019), Thailand (Piewbang et al., 2020), Malaysia (Anpuanandam et al., 2021), United Kingdom (Jeanes et al., 2022), United States (Stone et al., 2022), Hong Kong (Capozza et al., 2023), and Japan (Takahashi et al., 2022). DCH has a partially double-stranded, circular DNA genome of ∼3,200 bases in length (Aghazadeh et al., 2018). As observed with other *Orthohepadnavirus* species, the genome of DCH encodes polymerase, surface, core, and X proteins. However, the viral replication process of DHC is largely unknown, especially the entry into target cells.

HBV uses three envelope glycoproteins: Large (preS1 + preS2 + S), Middle (preS2 + S), and Small (S only) proteins for infection. Notably, the myristoylated preS1 domain is responsible for interaction with the cellular receptor (Engelke et al., 2006; Gripon et al., 2005; Le Seyec et al., 1999; Schulze et al., 2010). NTCP, a sodium-dependent bile acid symporter, is the major cell receptor for HBV (Yan et al., 2012). Therefore, considerable effort has been made to interrupt the interaction between preS1 and NTCP to develop an entry inhibitor. Myrcludex B, a synthetic lipopeptide derived from the preS1 domain of the HBV envelope protein, specifically targets hepatocytes to efficiently block de novo HBV infection (Gripon et al., 2005; Schulze et al., 2010), and has consequently received approval in Europe to treat patients.

In this study, we sought to use DCH as a model for HBV infection and investigated the entry pathway of DCH using the DCH-derived preS1 peptide. We demonstrated that the DCH-derived preS1 peptide efficiently binds to cat, and importantly, human NTCP. Consistent with findings with HBV (Takeuchi et al., 2019), the G158 residue of NTCP determines the species specificity of DCH, suggesting that the binding mode of preS1 and NTCP is conserved between HBV and DCH and that there is a potential risk of inter-species transmission of DCH. Myrcludex B potently inhibited binding of the DCH-derived preS1 peptide to human and cat NTCPs, suggesting that mode of inhibition can be evaluated in DCH-infected cats. Overall, this study provides a promising strategy for establishing a novel animal model for HBV infection, which could help develop therapeutic measures in HBV-infected individuals.

## 2. Materials and Methods

### 2.1. Plasmids

NTCP cDNA sequences of 19 animal species (**Table. 1**) were synthesized (Twist Bioscience) with codon optimization for expression in human cells. Synthesized DNA sequences are summarized in **Supplementary Table 1**. For transient expression of NTCP, inserts encoding NTCP cDNA were cloned into a pCAGGS vector (Niwa et al., 1991) predigested with EcoRI-HF (NEB, Cat# R3101M) and NheI-HF (NEB, Cat# R3131M) using In-Fusion Snap Assembly Master Mix (TaKaRa, Cat# Z8947N). In addition, for stable expression of NTCP, inserts encoding NTCP cDNA of human, cat, and cynomolgus monkey were cloned into a pLVSIN-CMV Hyg vector (TaKaRa, Cat# 6182) predigested with NotI-HF (NEB, Cat# R3189L) and BamHI-HF (NEB, Cat# R3136L) using In-Fusion Snap Assembly Master Mix. Plasmids were amplified using NEB 5-alpha F′Iq Competent *Escherichia coli* (High Efficiency) (NEB, Cat# C2992H), and isolated with PureYield Plasmid Miniprep System (Promega, Cat# A1222). Sequences of all plasmids were verified using a SupreDye v3.1 Cycle Sequencing Kit (M&S TechnoSystems, Cat# 063001) with a Spectrum Compact CE System (Promega). The psPAX2-IN/HiBiT plasmid was a kind gift from Dr. Kenzo Tokunaga (Ozono et al., 2020).

**Table 1.** NTCP amino acid sequence of 19 animal species tested in this study.

| Species | Common name | GenBank Acc. No |
| --- | --- | --- |
| *Homo sapiens* | Human | NP 003040.1 |
| *Macaca fascicularis* | Cynomolgus monkey | XP 045252552.1 |
| *Saimiri sciureus* | Squirrel monkey | ALX72398.1 |
| *Aotus nancymae* | Nancy Ma's night monkey | XP 012293622.1 |
| *Felis catus* | Cat | XP 003987831.1 |
| *Panthera tigris* | Tiger | XP 007095203.2 |
| *Leopardus geoffroyi* | Geoffroyi's cat | XP 045304082.1 |
| *Canis lupus familiaris* | Dog | XP 537494.2 |
| *Tupaia belangeri* | Treeshrew | AFK93890.1 |
| *Rhinolopus ferrumequinum* | Horseshoe bat | KAF6352409.1 |
| *Pteropus alecto* | Black fruit bat | XP 006915243.1 |
| *Sus scrofa* | Pig | XP 001927730.1 |
| *Bos taurus* | Cattle | NP 001039804.1 |
| *Tursiops truncatus* | Dolphin | XP 033706895.1 |
| *Marmota monax* | Woodchuck | XP 046316751.1 |
| *Incidomys tridecemlineatus* | Ground squirrel | XP 005322900.1 |
| *Mus musculus* | Mouse | NP 001171032.1 |
| *Rattus norvegicus* | Rat | NP 058743.1 |
| *Ornithorhynchus anatinus* | Platypus | XP 028932136.1 |

### 2.2. Generation of plasmids encoding NTCPs mutants

To generate NTCP mutants, mutagenesis was performed via overlapping PCR using PrimeSTAR GXL DNA polymerase (TaKaRa, Cat# R050A) with the primers listed in **Supplementary Table 2**. The PCR protocol consisted of 35 cycles at 98°C for 10 s, 60°C for 15 s, and 68°C for 1 min, followed by 68°C for 7 min. Amplified PCR fragments encoding the intended mutation were cloned into a pCAGGS vector as described above. Plasmids were verified by sequencing.

### 2.3. Cell culture

Lenti-X 293T cells (TaKaRa, Cat# Z2180N) were cultured in Dulbecco’s modified Eagle medium (DMEM) (Nacalai Tesque, Cat# 08458-16) supplemented with 10% fetal bovine serum and 1× penicillin–streptomycin (Nacalai Tesque, Cat# 09367-34).

### 2.4. Generation of Lenti-X 293T cells stably expressing NTCP molecules

To prepare an HIV-1-based lentiviral vector, Lenti-X 293T cells were cotransfected with plasmids psPAX2-IN/HiBiT, pLVSIN-CMV Hyg-NTCP, and pMD2.G using TransIT-293 Transfection Reagent (TaKaRa, Cat# V2700) in Opti-MEM I Reduced Serum Medium (Thermo Fisher Scientific, Cat# 31985062). The supernatant was filtered 2 days after transfection, and collected lentiviral vectors were used to infect Lenti-X 293T cells. Cells were cultured with 500 μg/mL Hygromycin B (Nacalai Tesque, Cat# 09287-84) for 6 days. Then, single-cell cloning was performed, and the expression level of NTCP in each clone was evaluated by western blotting.

### 2.5. preS1 peptide binding assay

Lipopeptide myr-47WT, a N-terminally myristoylated peptide comprising residues 2 to 47 of the preS1 region of HBV, and myr-47N9K, where Asn at position 9 is substituted with Lys, were synthesized by Scrum Co. (Tokyo, Japan). In addition, lipopeptide myr-47-DCH-WT, an N-terminal myristoylated peptide comprising residues 2 to 47 of the preS1 region of DCH, and myr-47-DCH-N9K were synthesized.

Lenti-X 293T cells transiently expressing NTCP (3 × 10^6^) were plated in a Celltight C-1 Plate 96F plate (Wako, Cat# 631-26891). After overnight incubation, cells were cotransfected with pCAGGS-NTCP plasmids together with pCAGGS-mCherry2 plasmid using TransIT-293 Transfection Reagent in Opti-MEM I Reduced Serum Medium to normalize the transfection efficiency. At 24 h after transfection, cells were inoculated with the corresponding peptides and incubated at 37°C for 60 min. After two washes with Gibco William’s E Medium (Thermo Fisher Scientific, Cat# A1217601), cells were observed under the EVOS M7000 Imaging System (Thermo Fisher Scientific). The cells were also analyzed via flow cytometry. The mCherry2-positive population was gated, and the percentage of FAM-positive cells were analyzed using FlowJo v10.8.1 (Becton, Dickinson and Company).

### 2.6. Inhibition of preS1 peptide binding by Myrcludex B treatment

Cells were prepared as described above and were then treated with 1000, 100, 10, 1, 0.1, 0.01, or 0.001 nM Myrcludex B (Selleck, Cat# P1105) at 37°C for 1 h. After two washes with DMEM, cells were inoculated with FAM-labeled preS1 peptides and incubated at 37°C for 1 h. The binding of FAM-labeled preS1 peptides was measured as described above. In addition, the cytotoxicity of Myrcludex B was evaluated with the Cell Counting Kit-8 (Dojindo, Cat# CK04) according to the manufacturer’s instruction.

### 2.7. Western blotting

To check the expression level of NTCPs, pelleted cells were lysed in 2× Bolt LDS sample buffer (Thermo Fisher Scientific, Cat# B0008) containing 2% β-mercaptoethanol (Bio-Rad, Cat# 1610710) and incubated at 70°C for 10 min. Expression of HA-tagged NTCPs in Lenti-X 293T cells was evaluated using SimpleWestern Abby (ProteinSimple) with an anti-HA Tag (6E2) mouse monoclonal antibody (CST, Cat# 2367S, ×250) and an Anti-Mouse Detection Module (ProteinSimple, Cat# DM-002). In addition, the amount of input protein was measured using a Total Protein Detection Module (ProteinSimple, Cat# DM-TP01).

### 2.8. Alignment of the regions around residue 158 of NTCPs and phylogenetic analysis

The regions around residue 158 of the NTCPs from 20 animal species were aligned using the MUSCLE algorithm in MEGA X (MEGA Software). The parameters of alignment were: gap open, −2.90; gap extend, 0.00; and hydrophobicity multiplier, 1.20.

A phylogenetic tree was built using alignment of NTCP amino acid sequences retrieved from public databases and the evolutionary analysis were conducted in MEGA X (Kumar et al., 2018; Stecher et al., 2020). The evolutionary history was inferred using the Maximum Likelihood method and JTT matrix-based mode (Jones et al., 1992). Initial tree for the heuristic search were obtained by applying Neighbor-Joining method to a matrix of pairwise distances estimated using JJT model. A discrete Gamma distribution was used to model evolutionary rate differences among sites. The tree is drawn to scale, with branch lengths measured in the number of substitutions per site.

### 2.9. Calculation of identity of NTCP among animal species

The identity of NTCPs among animal species was calculated using MEGA X (Kumar et al., 2018; Stecher et al., 2020) with a pairwise distance matrix. Analyses were conducted using the Poisson correction model. The rate variation among sites was modeled with a gamma distribution (shape parameter = 5). All ambiguous positions were removed for each sequence pair (pairwise deletion option).

### 2.10. Statistical analysis

Differences in binding between wild type and amino acid 158 mutants were evaluated using unpaired, two-tailed Student’s *t*-test. A *p* ≤ 0.05 was considered statistically significant. Analysis was performed using Prism 9 software v9.1.1 (GraphPad Software).

## 3. Results

### 3.1. Genetic similarity of preS1 between DCH and HBV

We aligned the preS1 sequence of 13 strains of DCH, five strains of HBV (genotype A, B, C, D, and F), two strains of Arctic ground squirrel hepatitis B virus, one strain of woodchuck hepatitis virus and those of other viruses belonging to *Orthohepadnavirus* (**Figure 1A**). As expected, HBV has different clades based on the genotype (**Figure 1B**), and DCH was also distributed into several clades. Notably, the DCH preS1 sequence is genetically closer to that of HBV rather than that of woodchuck hepatitis virus or Arctic ground squirrel hepatitis B virus, suggesting that DCH preS1 may exhibit a similar function to that of HBV preS1.

**Figure 1.**
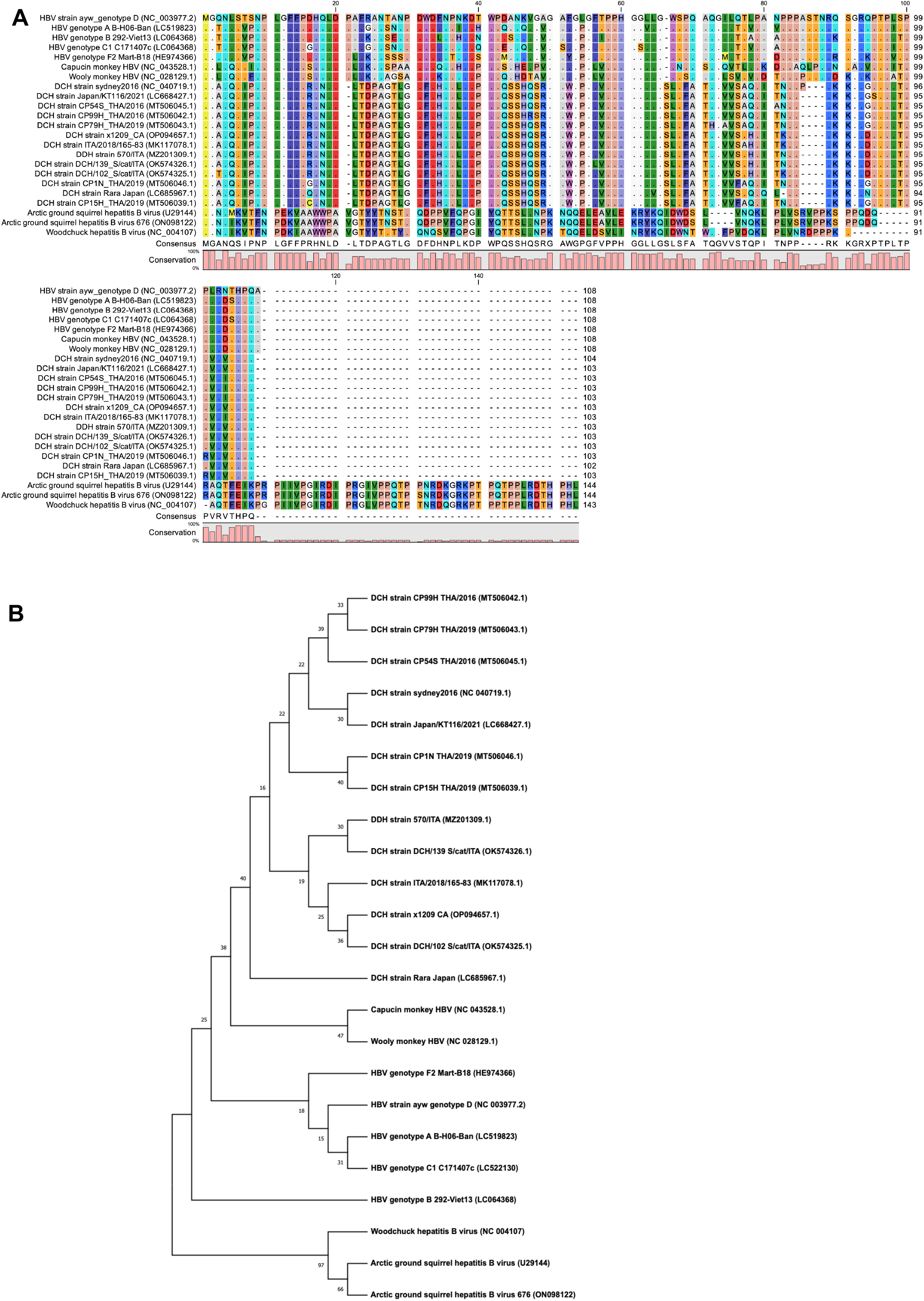
Sequence alignment and phylogenetic analysis of the preS1 amino acids of orthohepadnaviruses. (A) Amino acid sequences of the preS1 domain of DCH aligned with the corresponding regions of other *Orthohepadnavirus* species. Residue numbering was based on HBV genotype D strain. (B) A phylogenetic tree built using alignment of preS1 amino acid sequences derived from orthohepadnaviruses retrieved from public databases. The evolutionary history was inferred using the Neighbor-Joining method. The percentage of replicate trees where the associated taxa clustered together in the bootstrap test (1,000 replicates) are shown next to the branches.

### 3.2. Genetic similarity of NTCPs among mammals

We next analyzed the NTCP genetic similarities between humans and other animals, including cat (**Figure 2**). The phylogenetic tree shows that the feline NTCP is genetically close to primates despite being in a different branch. The similarity rates of human and cat were 86% and 82% at the nucleotide and amino acid levels, respectively, based on the estimation of evolutionary differences between sequences (**Supplementary Table 2**). Higher sequence distances were seen for NTCPs from other mammal species, such as dolphin, woodchuck, ground squirrel, mouse, rat, and platypus (sequence identities of <80% compared with human NTCP). Nonetheless, some mammals with a higher NTCP distance from humans contained the same amino acid at position 158 (**Figure 2**).

**Figure 2.**
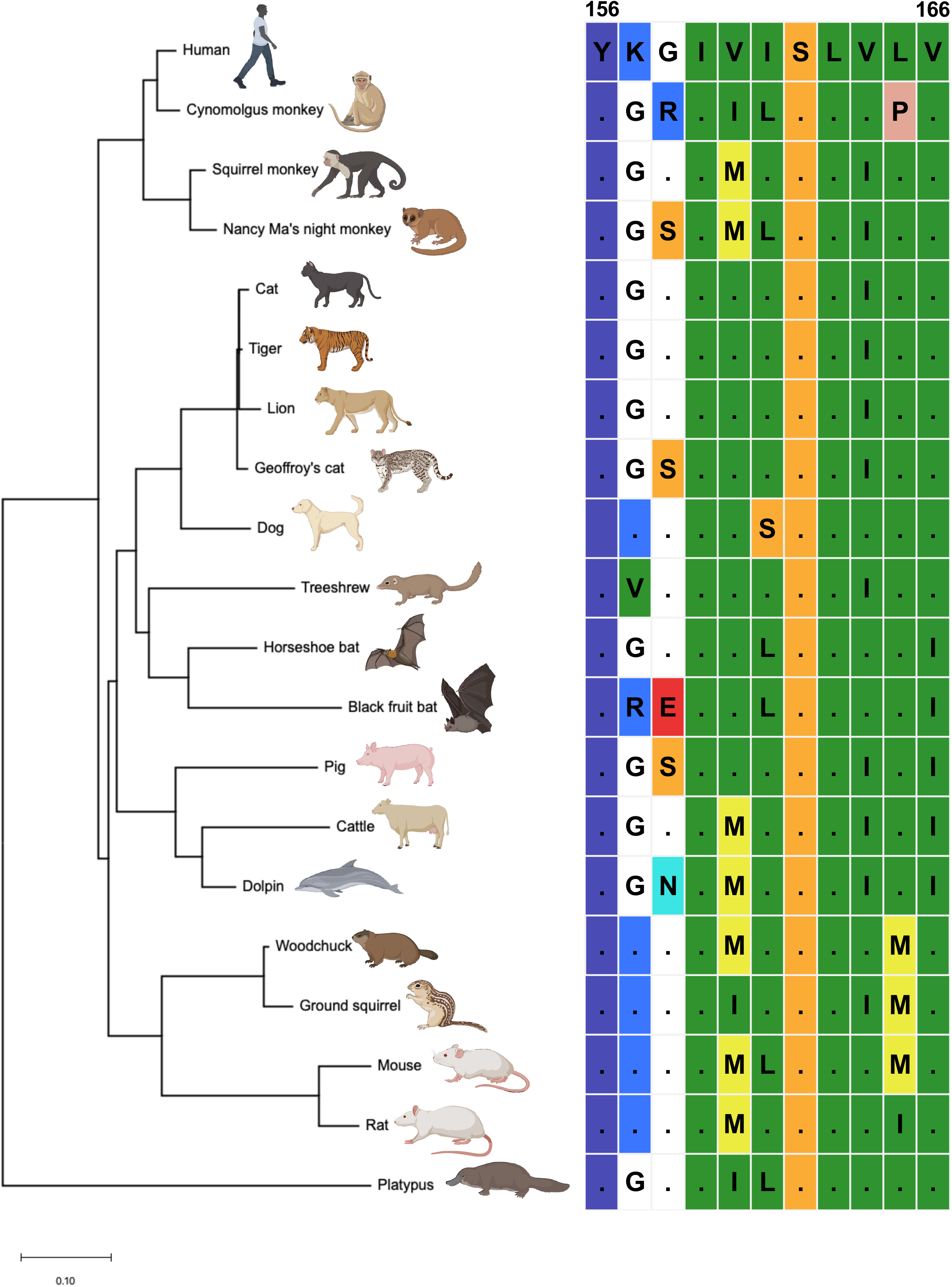
Sequence alignment of mammalian NTCP (residue 156–166) and phylogenetic analysis of the entire mammalian NTCP sequence. Mammalian NTCP amino acids (156–166) were aligned and the phylogenetic tree of the entire mammalian NTCP sequence was constructed. The residue numbering was based on the human NTCP sequence as a reference.

### 3.3. The DCH preS1 peptide binds to both cat and human NTCPs

Myristoylated HBV preS1 peptide (47 amino acids in length) can be used to test the binding of preS1 to its counterpart NTCP (Schulze et al., 2010). Therefore, we synthesized HBV and DCH FAM-labeled, myristoylated preS1 peptides. Lenti-X 293T cells transiently expressing either human NTCP (HumanNTCP) or cat NTCP (CatNTCP) were tested for binding to myristoylated preS1 peptides. To distinguish between transfected and untransfected cells, we cotransfected a plasmid encoding mCherry2 fluorescent protein (**Figure 3A**) and measured the number of FAM-positive cells in the mCherry2-positive population, which should express each NTCP molecule, via fluorescent microscopy or flow cytometry (**Supplementary Figure 1**). The expression level of HA-tagged NTCP molecules was confirmed with western blotting (**Figure 3B**). We observed marginal binding of preS1 peptides to cells transfected with empty vector (**Figure 3C**). In contrast, the HBV-derived preS1 peptide efficiently bound to cells expressing HumanNTCP (**Figures 3C and 3D**); binding was not observed with HBV-derived preS1 peptide harboring the N9K substitution (**Figure 3D**), which is consistent with previous studies demonstrating that the N9 residue is critical for interaction with HumanNTCP (Yan et al., 2012). Moreover, the HBV-derived preS1 peptide showed significant binding to CatNTCP to a level similar to that of HumanNTCP. As previously described (Takeuchi et al., 2019), cynomolgus monkey NTCP (CmNTCP) did not support binding of the HBV-derived preS1 peptide. Furthermore, we found DCH-derived preS1 peptide efficiently bound to both CatNTCP and HumanNTCP but not to CmNTCP (**Figures 3C and 3D**), which was confirmed with transiently transfected cells and cells that stably expressed the corresponding NTCP molecules (**Supplementary Figures 2A and 2B**). Furthermore, similar to HBV preS1 peptide, the DCH-derived preS1 peptide harboring the N9K substitution failed to bind any NTCPs (**Figure 3D**).

**Figure 3.**
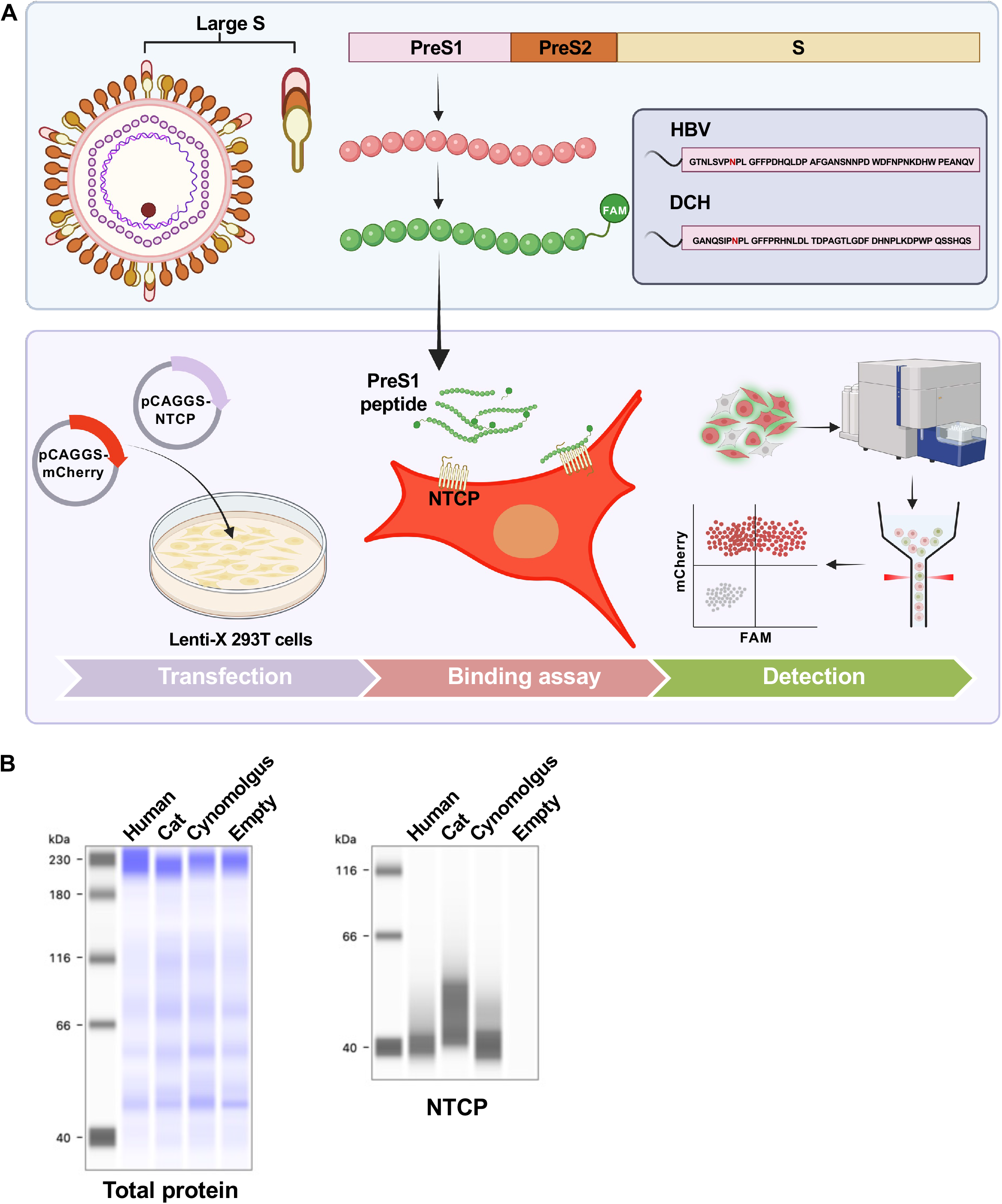
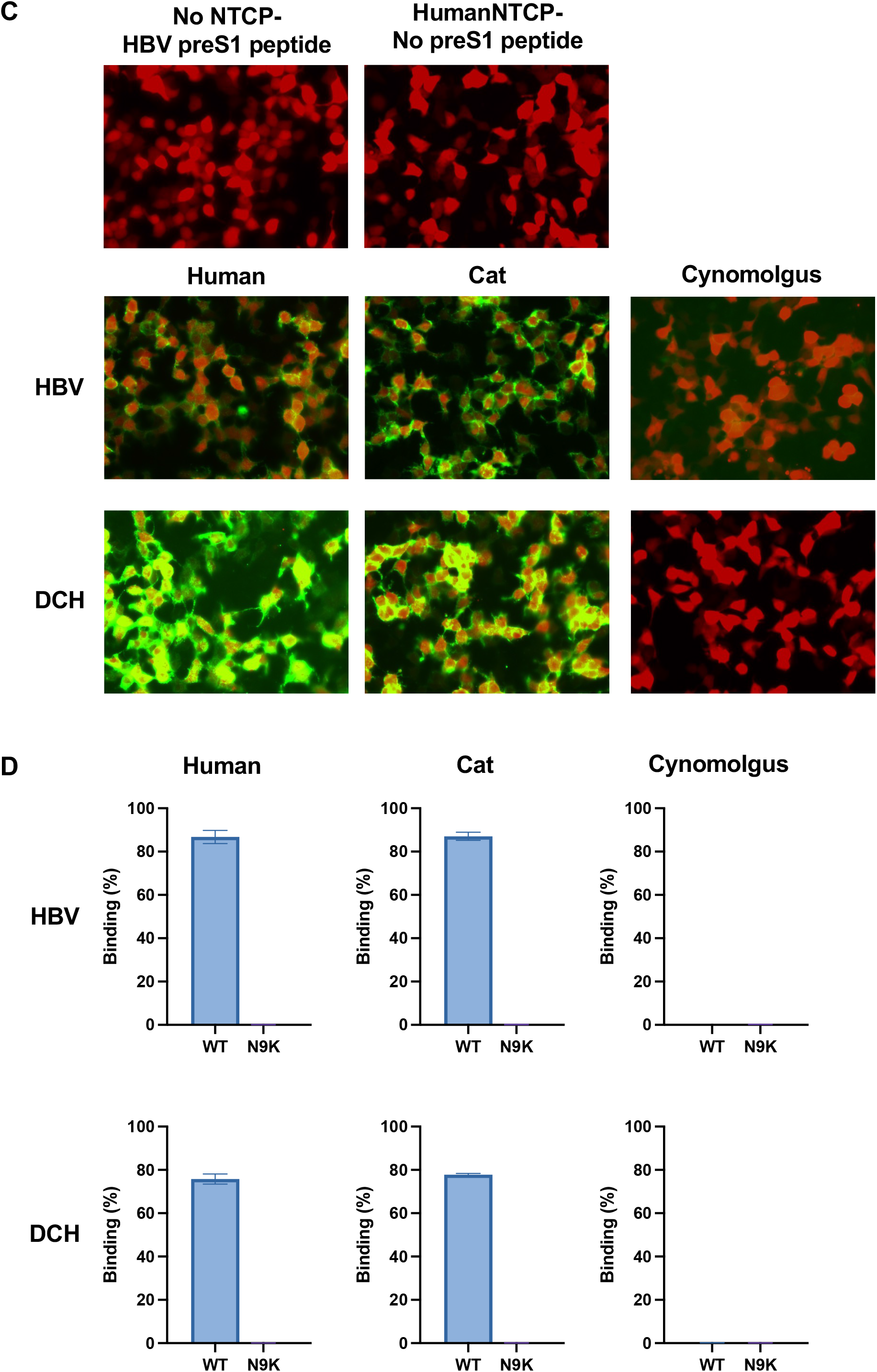
DCH preS1 peptide binds to both CatNTCP and HumanNTCP. (A) Schematic overview of the binding assay of FAM-labeled preS1 peptides to Lenti-X 293T cells cotransfected with human, cat, or cynomolgus monkey NTCPs and mCherry-expressing plasmid. For analyses with a fluorescent microscopy or a flow cytometer, the presence of FAM-positive cells in the mCherry2-positive population was analyzed. (B) Expression levels of human, cat, or cynomolgus monkey NTCPs in transiently transfected Lenti-X 293T cells was examined via western blotting (C) Cells were inoculated with HBV or DCH-derived preS1 peptides harboring wild-type amino acids or N9K substitution. HumanNTCP-expressing cells without the HBV-derived preS1 peptide and normal cells with the HBV-derived preS1 peptide were negative controls. (D) Binding of HBV or DCH-derived preS1 peptides to NTCPs was measured via flow cytometry.

To better understand the species tropism of the DCH preS1 peptide, we expanded our analysis to NTCP molecules from other mammals, including squirrel monkey, Nancy Ma’s night monkey, tiger, Geoffroy’s cat, dog, horseshoe bat, black fruit bat, pig, cow, dolphin, woodchuck, ground squirrel, mouse, rat, and platypus (**Figures 2 and 4A**). Notably, squirrel monkey, horseshoe bat, ground squirrel, and woodchuck are natural hosts for *Orthohepadnavirus* species (Magnius et al., 2020). The DCH-derived preS1 peptide bound to NTCP from a wide range of mammals (**Figures 4B and 4C**). Although possible transmission cases of DCH to dogs have been reported (Choi et al., 2022; Diakoudi et al., 2022), the binding of the DCH-derived preS1 peptide to dog NTCP was weak.

**Figure 4.**
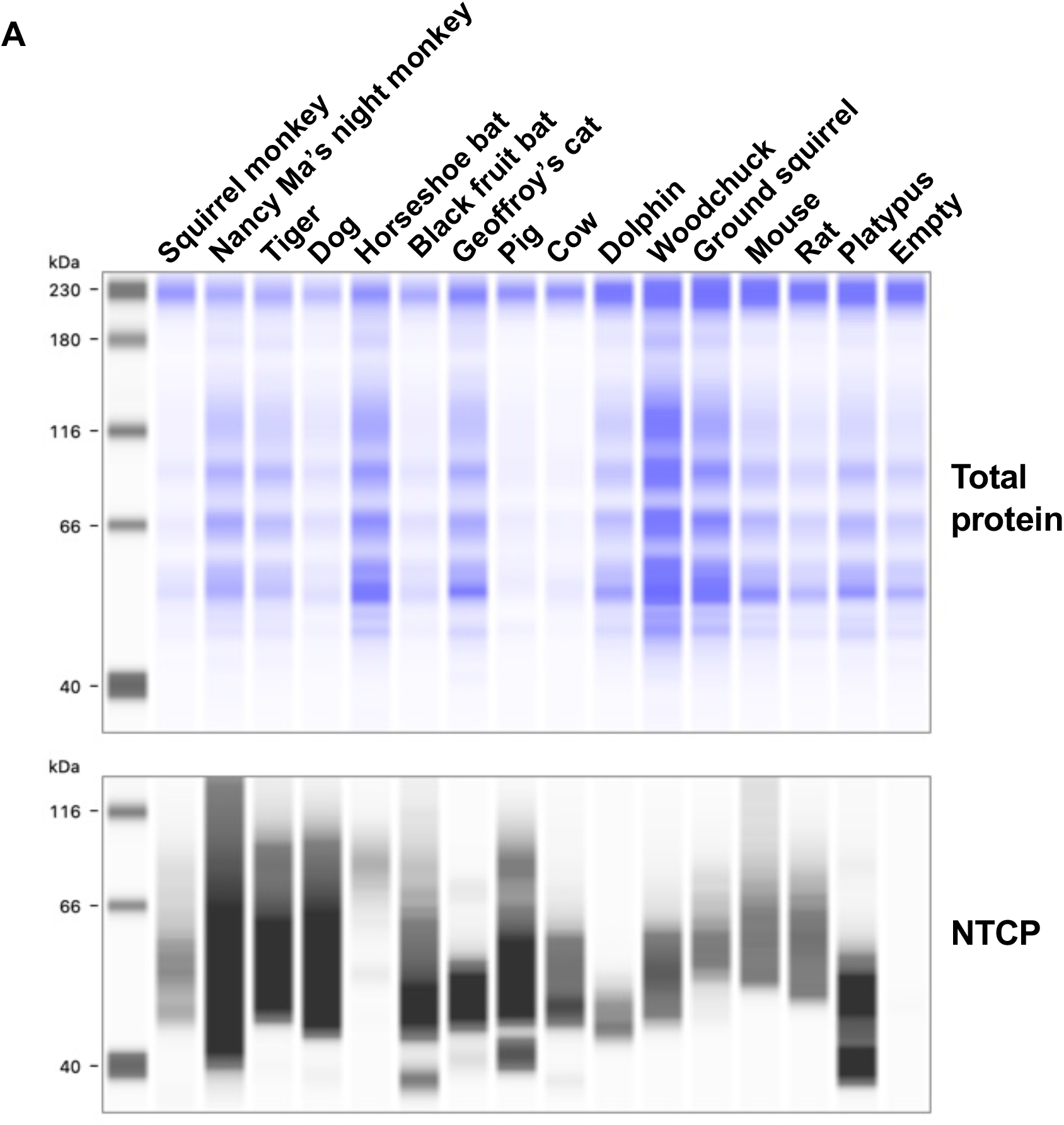
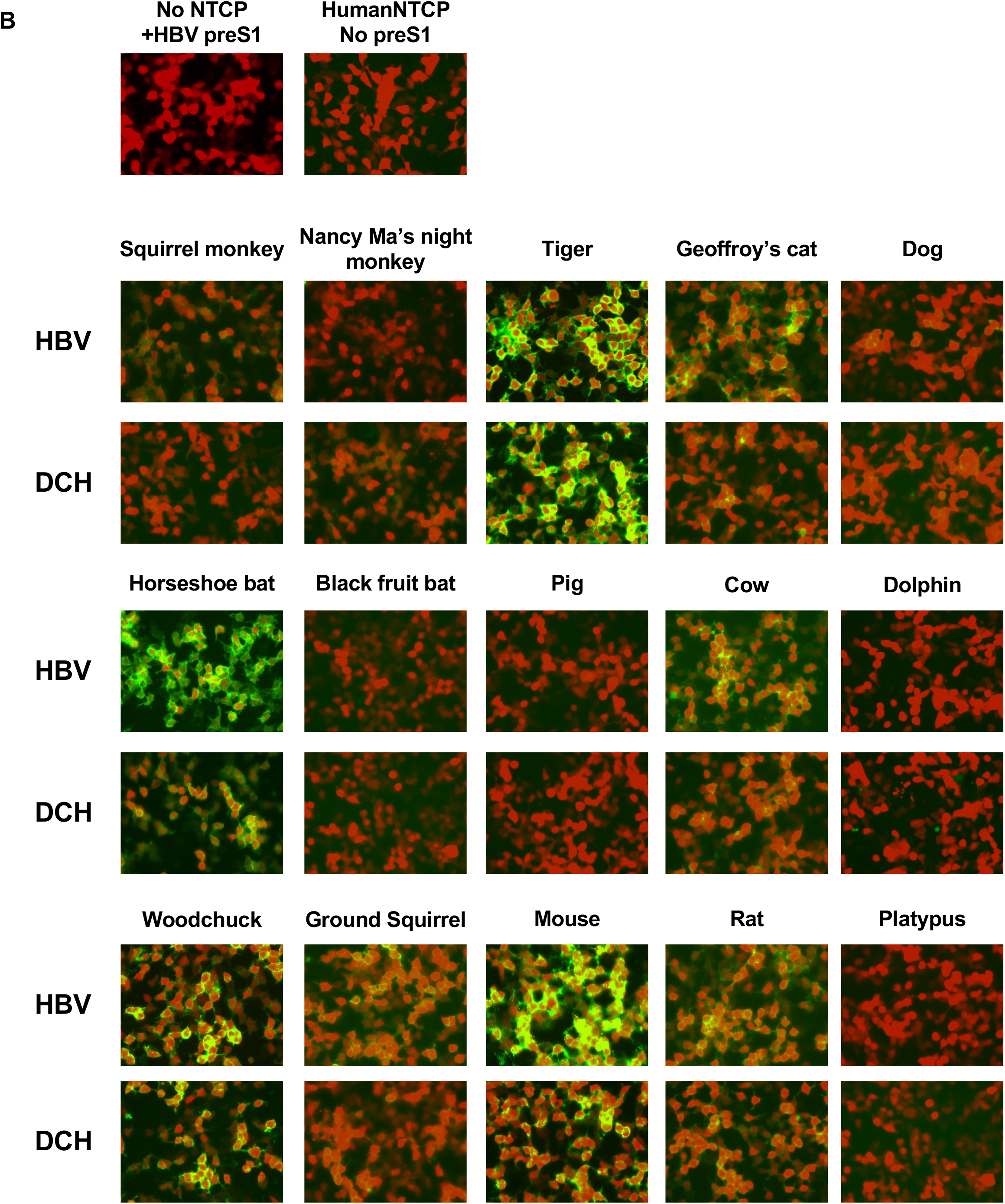
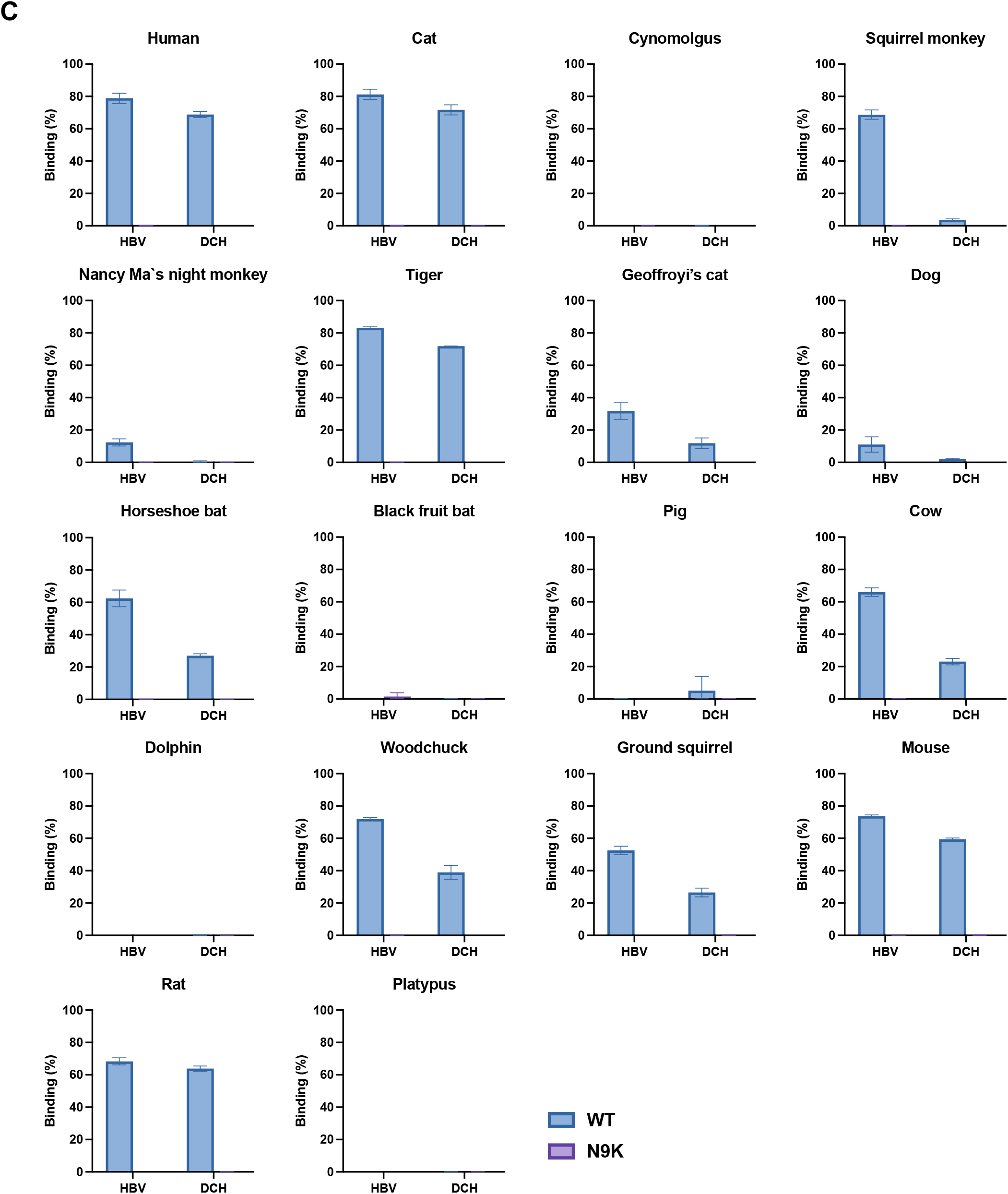
Binding spectrum of the DCH-derived preS1 to mammalian NTCPs. (A) Expression levels of mammalian NTCPs in transiently transfected Lenti-X 293T cells examined via western blotting. (B) Cells were inoculated with HBV or DCH-derived preS1 peptides harboring wild-type amino acids or N9K substitution. HumanNTCP-expressing cells without the HBV-derived preS1 peptide and normal cells with the HBV-derived preS1 peptide were negative controls. (C) Binding of HBV or DCH-derived preS1 peptides to the NTCPs measured via flow cytometry.

### 3.4. The G158 residue of NTCP determines species specificity of the DCH preS1 peptide binding

Previous studies demonstrated that G158 of HumanNTCP is critical for interaction with the HBV-derived preS1 peptide (Takeuchi et al., 2019). To evaluate whether this applies to the DCH-derived preS1 peptide, we generated mutant NTCPs including HumanNTCP (G158R), CatNTCP (G158R), and CmNTCP (R158G) (**Figure 5A**). The HBV-derived preS1 peptide lost binding to HumanNTCP (G158R) and CatNTCP (G158R) but acquired significant binding to CmNTCP (R158G) (**Figures 5B and 5C**). Similarly, the DCH-derived preS1 peptide lost binding to HumanNTCP (G158R) and CatNTCP (G158R) but showed significant binding to CmNTCP (R158G) (**Figures 5B and 5C**), suggesting that the binding pattern of the DCH-derived preS1 peptide is similar to that of the HBV-derived preS1 peptide. The amino acid at position 158 is therefore critical for binding with the DCH-derived preS1 peptide.

**Figure 5.**
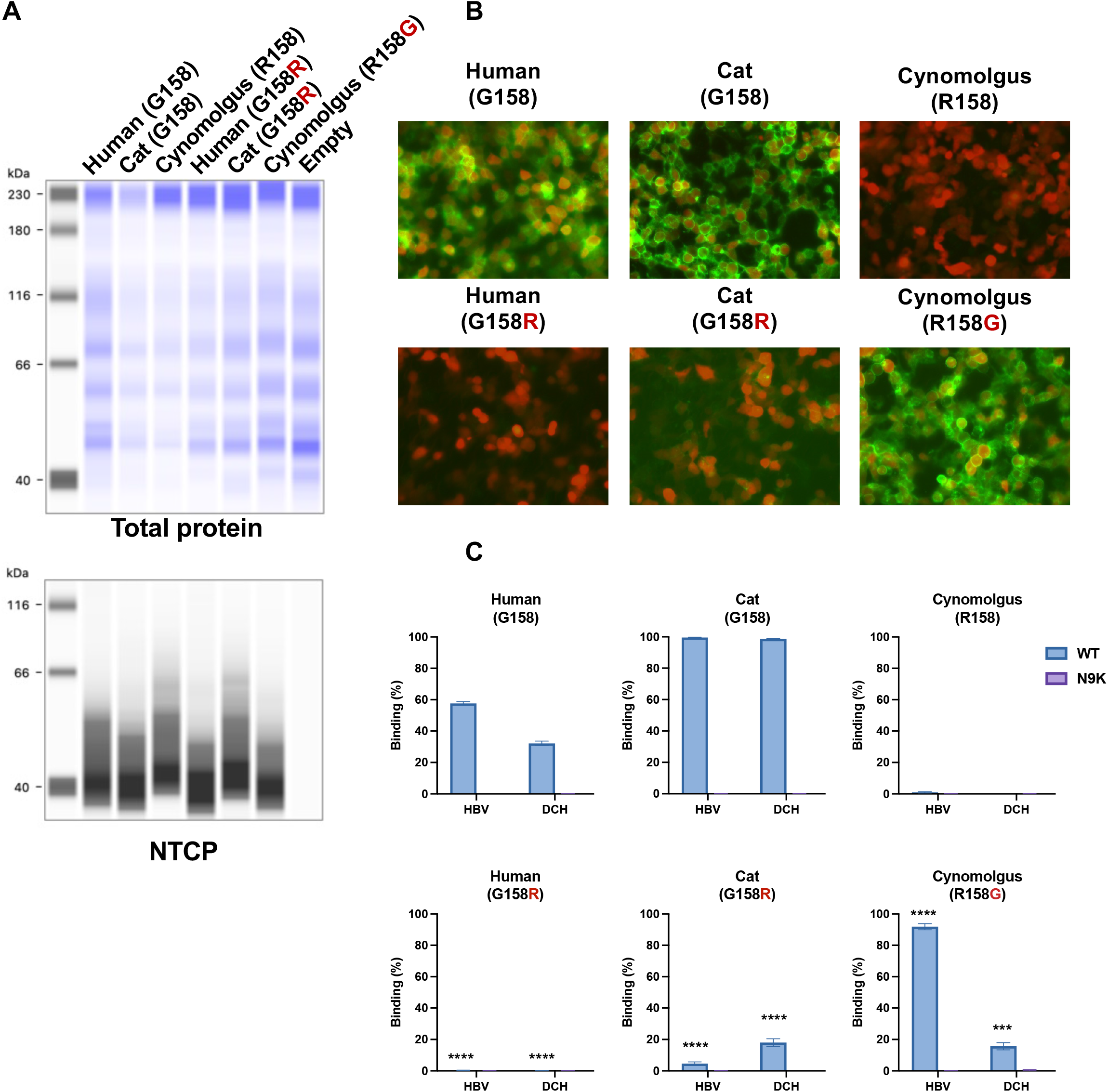
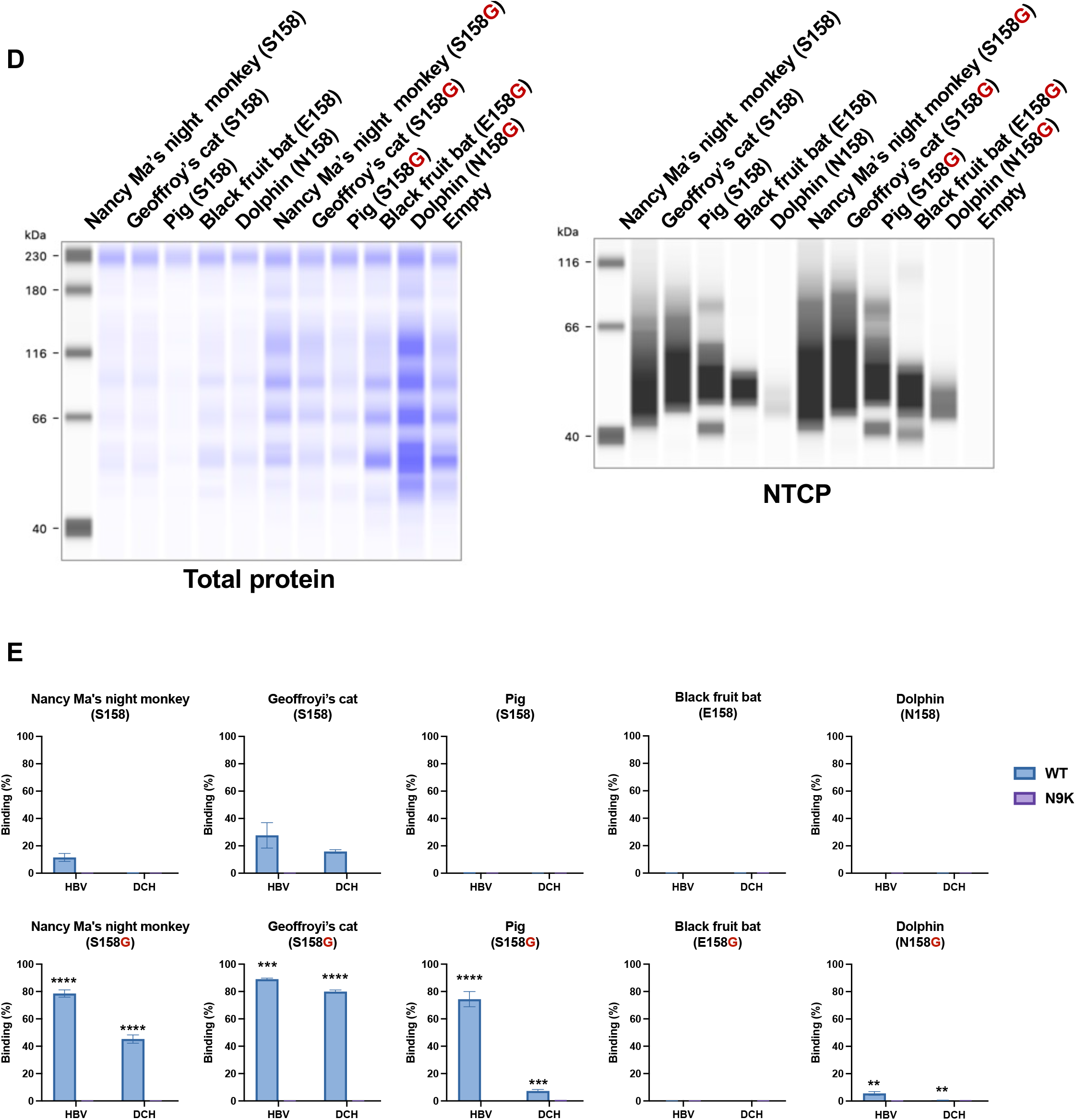
The G158 residue of NTCP determines species specificity of the DCH preS1 peptide binding. (A) Expression levels of human (G158, WT; G158R), Cat (G158, WT; G158R), cynomolgus monkey (R158, WT; R158G) NTCPs in transiently transfected Lenti-X 293T cells examined via western blotting (B) Cells were inoculated with HBV or DCH-derived preS1 peptides harboring wild-type amino acids or N9K substitution. HumanNTCP-expressing cells without the HBV-derived preS1 peptide and normal cells with the HBV-derived preS1 peptide were negative controls. (C) Binding of HBV or DCH-derived preS1 peptides to the NTCPs measured via flow cytometry. The results are presented as the mean and standard deviation of triplicate measurements from one assay, and they are representative of at least three independent experiments. Differences in binding between wild type and amino acid 158 mutants were evaluated using unpaired, two-tailed Student’s *t*-test. \*\*\*\**p* < 0.0001, \*\*\**p* < 0.001. (D) Expression levels of Nancy Ma’s night monkey (S158, WT; S158G), Geoffroy’s cat (S158, WT; S158G), pig (S158, WT; S158G), black fruit bat (E158, WT; E158G) NTCPs in transiently transfected Lenti-X 293T cells examined via western blotting. (E) Cells were inoculated with the HBV or DCH-derived preS1 peptides harboring wild-type amino acids or N9K substitution. HumanNTCP-expressing cells without the HBV-derived preS1 peptide and normal cells with the HBV-derived preS1 peptide were negative controls. The results are presented as the mean and standard deviation of triplicate measurements from one assay, and they are representative of at least three independent experiments. Differences in binding between wild type and amino acid 158 mutants were evaluated using unpaired, two-tailed Student’s *t*-test. \*\*\*\**p* < 0.0001, \*\*\**p* < 0.001, \*\**p* < 0.01.

In addition to cynomolgus monkey (R158), non-G residues at this position can be found in animals such as Nancy Ma’s night monkey (S158), Geoffroy’s cat (S158), black fruit bat (E158), pig (S158), and dolphin (N158) (**Figure 2**). Binding analyses demonstrated that these NTCPs do not support binding to either the HBV-derived or the DCH-derived preS1 peptides (**Figures 4B and 4C**). To test whether changing the residue to glycine affects the binding of the DCH-derived preS1 peptide, we performed a mutagenesis experiment to introduce G158 to each of these NTCPs (**Figure 5D**). Substitution of residue 158 with glycine significantly augmented binding of these NTCPs to the HBV-derived and the DCH-derived preS1 peptides (**Figure 5E**). These findings demonstrate that a single amino acid at position 158 of NTCP determines the interaction with preS1 peptide and that this dependency is shared by HBV and DCH.

### 3.5. Myrcludex B treatment blocks binding of the DCH preS1 peptide to both human and cat NTCPs

Myrcludex B specifically inhibits the interaction between HBV-derived preS1 and HumanNTCP (Gripon et al., 2005; Petersen et al., 2008). Therefore, we investigated the effect of Myrcludex B on the interaction between the DCH-derived preS1 peptide and HumanNTCP or CatNTCP (**Figure 6A**). As previously shown (Gripon et al., 2005; Petersen et al., 2008), Myrcludex B efficiently blocked the interaction between the HBV-derived preS1 peptide and HumanNTCP (IC_50_: 57.55 nM) (**Figure 6B**) without any cytotoxicity (**Supplementary Figure 3**). A comparable effect was observed with a combination of HBV-derived preS1 peptide and CatNTCP (IC_50_: 54.3 nM), suggesting that HumanNTCP and CatNTCP share the same mode of action for Myrcludex B-mediated inhibition. Notably, Myrcludex B efficiently blocked interaction between the DCH-derived preS1 peptide and HumanNTCP or CatNTCP at a similar concentration (IC_50_: 33.79 and 35.55 nM, respectively) (**Figure 6B**).

**Figure 6.**
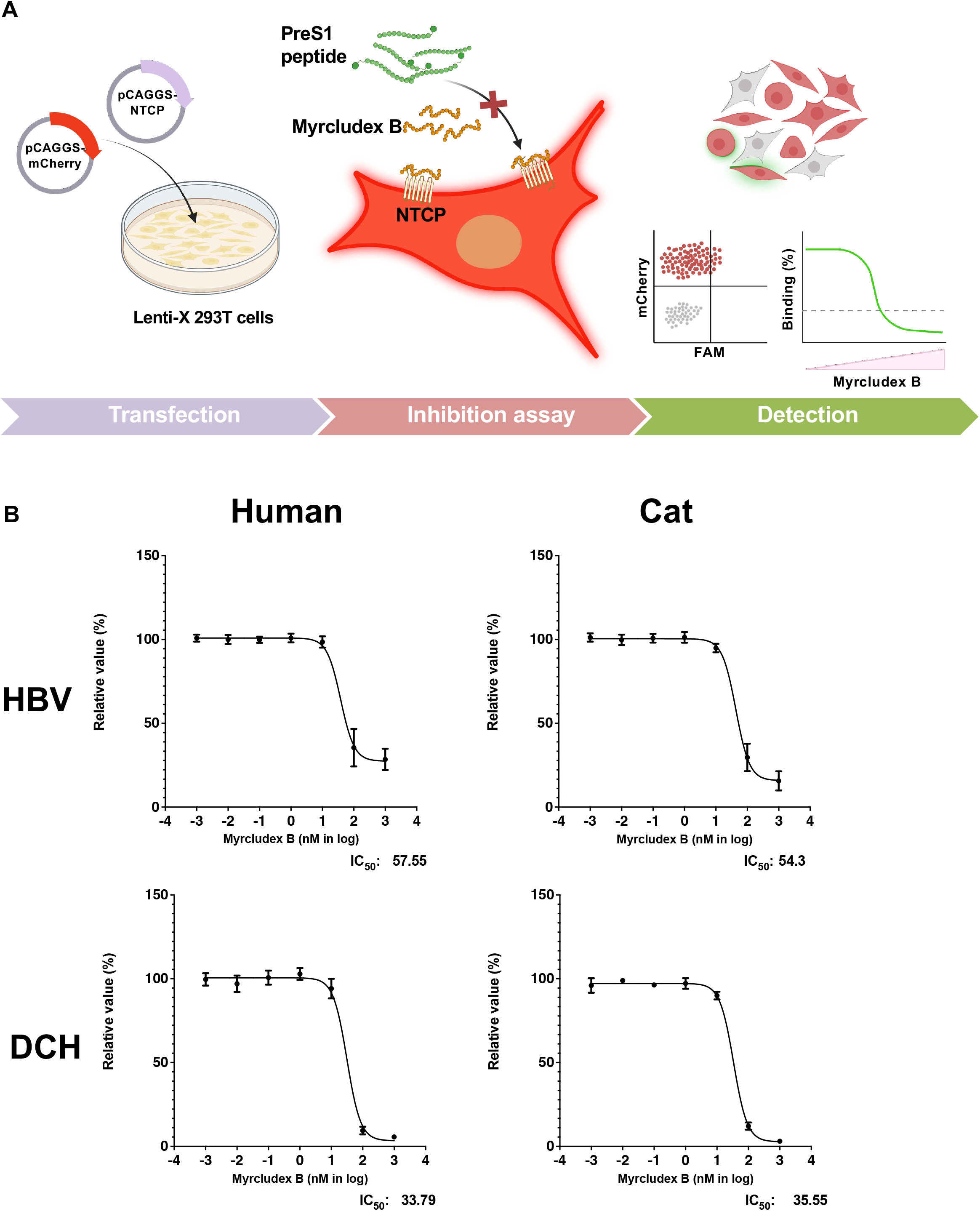
Myrcludex B treatment blocks binding of the DCH preS1 peptide to both human and cat NTCPs. (A) Schematic overview of the Myrcludex B-mediated inhibition assay using FAM-labeled PreS1 peptides to Lenti-X 293T cells cotransfected with human or cat NTCPs and a mCherry-expressing plasmid. For analyses with a fluorescent microscopy or a flow cytometer, the presence of FAM-positive cells in the mCherry2-positive population was analyzed. (B) Myrcludex B inhibits the binding of HBV or DCH-derived preS1 peptides to human or cat NTCPs; IC_50_ values were calculated.

Collectively, the HBV-derived preS1 peptide and the DCH-derived preS1 peptide share the susceptibility to Myrcludex B, suggesting that these viruses depend on similar molecular interactions for binding to NTCP on the cellular surface.

## 4. Discussion

Herein, we demonstrated a significant similarity in cellular entry receptors between HBV and DCH. Both viruses utilize NTCP as a receptor, and more importantly, the HBV-derived preS1 peptide can bind to cat NTCP while the DCH-derived preS1 peptide can bind to human NTCP. Furthermore, we demonstrated that the DCH-derived preS1 peptide binds to NTCPs derived from a broad range of animal species, suggesting that DCH can be transmitted to these animals. Finally, we demonstrated that Myrcludex B inhibits interaction of the DCH-derived preS1 peptide with human and cat NTCPs, suggesting that DCH can be used for evaluation of entry inhibitors targeting HBV.

The phylogenetic tree of the preS1 sequence of *Orthohepadnavirus* species suggests that several strains of DCH are genetically closer to HBV than orthohepadnaviruses associated with other host species. This suggests that feline DCH infection can be a promising animal model for HBV infection. However, infectious DCH has yet to be isolated, and this should be a focus of continued research, specifically in cats. It is critical to understand whether pathogenesis of DCH in infected cats causes chronic hepatitis and hepatocellular carcinoma as HBV infection can do in humans. Furthermore, although an analysis based on a naturally infected cat suggested a low possibility of sexual transmission of DCH, this should be evaluated under controlled experimental conditions.

We identified CatNTCP as a binding molecule for the preS1 peptide of DCH (**Figures 3C and 3D**), suggesting that CatNTCP is a cellular receptor for DCH. Furthermore, mutagenesis of glycine at NTCP position 158 to arginine caused a loss in binding to the DCH-derived preS1 peptide for both HumanNTCP (G158R) or CatNTCP (G158R) but produced significant binding to CmNTCP (R158G) (**Figures 5B and 5C**), indicating the shared nature of the binding of DCH-and HBV-derived preS1 peptide to the NTCP receptors. Furthermore, DCH may be zoonotic in some circumstances as the DCH-derived preS1 peptide efficiently bound to NTCPs from a broad animal species, including humans (**Figure 3D and 4D**). However, infectious DCH culture is required to enable testing of DCH replication in human cells. Interestingly, DogNTCP did not support binding of the HBV-derived and the DCH-derived preS1 peptides (**Figure 3D**). Although DCH has been identified in dogs (Diakoudi et al., 2022), the low level of expression of DogNTCP in our system might contribute to this discrepancy and should be evaluated in future experiments.

Myrcludex B efficiently inhibited interaction of the DCH-derived preS1 peptide with CatNTCP (**Figure 6B**), suggesting that Myrcludex B can be used for treating cats chronically infected with DCH. Furthermore, clinical information from DCH-infected cats may provide insights for control of HBV infection and eliminating DCH from persistently infected cats could contribute to advances in HBV cure.

A previous study demonstrated that several amino acids of NTCP were under positive selection (Takeuchi et al., 2019), suggesting that NTCP has been a frontline for cell invasion by viruses, especially hepadnaviruses. Intriguingly, CmNTCP encodes R158 to provide escape from infection with HBV. We demonstrated that non-G residue were present at amino acid 158 of NTCP in several animal species (**Figure 2A**), and it would be interesting to investigate whether these species are natural hosts for *Orthohepadnavirus* and, if so, whether coevolution of their preS1 sequence has occurred to compensate for the decreased binding caused by the non-G residue at NTCP position 158. Another possible evolutionary tactic would be use of a different receptor molecule to infect the host in these species. We believe testing these possibilities will provide important insights on the arms race between hepadnaviruses and mammals.

The HBV-like virus has been identified in pigs (Vieira et al., 2014). However, the genetic distance between HBV and HBV-like virus in pig is very close, and this may be a possible contamination of viral DNA from HBV-infected humans. Our analysis demonstrated that the HBV-derived preS1 peptide did not bind to PigNTCP, supporting this hypothesis.

Alternatively, the HBV-like virus in pigs may carry other mutations in the preS1 domain for efficient binding with PigNTCP harboring S158. Considering a risk of food-borne transmission, whether pigs are a host for *Orthohepadnavirus* species should be investigated.

A limitation of this study is that we used synthesized lipopeptides to test the interaction between preS1 and the NTCP molecule and investigate the viral entry pathway. Therefore, the interaction between the DCH-derived preS1 peptide and NTCPs require experimental verification using infectious virus. Consequently, we are currently trying to isolate infectious DCH. We believe future study using infectious DCH will support our present findings.

In conclusion, we revealed a significant similarity in the cell entry pathway between HBV and DCH, suggesting a potential risk of inter-species transmission. Furthermore, our finding that Myrcludex B is active against both HBV and DCH suggests that the DCH infection in cats can be a promising animal model for HBV research.

## Glossary

DCH: Domestic cat hepadnavirus
HBV: Hepatitis B virus
MEGA: Molecular Evolutionary Genetics Analysis

## Acknowledgements

The authors thank Dr. Toru Okamoto (Osaka University) for insightful suggestions. The following reagents were obtained through the Addgene: pMD2.G was a gift from Dr. Didier Trono. The psPAX2-IN/HiBiT plasmid was a kind gift from Dr. Kenzo Tokunaga. The authors thank Ms. Tomoko Nishiuchi and Ms. Yuki Shibatani for their assistance. Figures 2, 3A and 6A were created with BioRender (https://biorender.com/).

## Author information

### Contributions

M.S. and A.S. designed experiments. M.S., A.O., and A.S. performed experiments. M.S. Y.K., and A.S. analyzed results. M.S. and A.S. wrote manuscript. All authors read and approve the manuscript.

### Data availability

Source data available on request.

## Ethics declarations

### Corresponding author

Akatsuki Saito.

