## Supplementary Materials for "Conserved use of the sodium/bile acid cotransporter (NTCP) as an entry receptor by hepatitis B virus and domestic cat hepadnavirus"

**Supplementary Figure 1. FACS gating strategy for measuring the preS1-polypetide binding to NTCPs.**

Lenti-X 293T cells cotransfected with plasmids encoding NTCP and mCherry were inoculated with HBV- or DCH-derived preS1 peptides. The presence of FAM-positive cells in the mCherry2-positive population was analyzed.

**Supplementary Figure 2. Binding of the preS1 peptides to Lenti-X 293T cells stably expressing NTCP molecules.**

Lenti-X 293T cells stably expressing human, cat, or cynomolgus monkey NTCPs were inoculated with HBV or DCH-derived preS1 peptides. The percentage of FAM-positive cells was analyzed.

**Supplementary Figure 3. CCK-8 cytotoxicity assay of Myrcludex B treatment**

Lenti-X 293T cells were treated with different concentrations of Myrcludex B. After 1 h incubation, the cell viability was measured using the CCK-8 cytotoxicity assay.

**Supplementary Table 1. Synthesized DNA for generating a plasmid expressing NTCP molecule**

| Species | Common name | Codon optimized DNA sequence |
|---|---|---|
| *Homo sapiens* | Human | ATGTACCCTTACGATGTACCTGACTACGCGGAGGCCCATAACGCGTC<br>TGCCCCGTTCAATTTTACCCTTCCCCCAAATTTCGGTAAAAGACCAAC<br>GGACCTGGCTCTCAGCGTCATATTGGTCTTCATGCTCTTTTTCATCAT<br>GCTTTCTTTGGGATGTACTATGGAGTTTTCCAAGATTAAAGCCCACCT<br>CTGGAAGCCCAAAGGGTTGGCTATAGCCCTTGTCGCACAATATGGAA<br>TCATGCCTCTGACCGCATTCGTGTTGGGGAAAGTTTTTCGGCTCAAAA<br>ACATCGAGGCTTTGGCCATTCTGGTCTGCGGGTGCAGTCCCGGCGGT<br>AACCTTTCAAACGTGTTCAGCCTTGCAATGAAAGGCGATATGAATCT<br>CTCAATAGTGATGACCACATGTAGCACGTTCTGTGCCCTTGGCATGAT<br>GCCACTGTTGCTTTATATCTATTCTAGAGGGATTTATGATGGAGACTT<br>GAAGGACAAGGTGCCCTACAAAGGTATCGTTATCTCACTCGTACTGG<br>TTTTGATCCCTTGTACAATAGGTATTGTGTTGAAATCAAAACGACCTC<br>AATACATGCGCTATGTTATCAAGGGCGGAATGATTATAATCCTTCTGT<br>GCTCTGTAGCGGTTACGGTTCTGTCTGCTATAAATGTTGGTAAAAGCA<br>TCATGTTTGCAATGACACCTCTGTTGATCGCTACATCATCCCTTATGC<br>CCTTTATTGGCTTTCTGCTTGGGTATGTACTGAGCGCACTCTTTTGCCT<br>CAATGGCCGGTGTCGGCGCACTGTGTCAATGGAAACTGGCTGTCAGA<br>ATGTCCAACTTTGCTCAACAATCTTGAATGTAGCTTTCCCTCCAGAGG<br>TCATCGGCCCCCTGTTCTTTTTTCCGCTTCTGTACATGATCTTCCAACT<br>CGGTGAGGGCTTGTTGTTGATTGCCATATTTTGGTGTTACGAGAAATT<br>CAAGACCCCTAAGGACAAGACAAAGATGATCTATACTGCCGCAACA<br>ACTGAGGAAACAATACCTGGGGCGCTCGGAAATGGCACATACAAGG<br>GAGAGGACTGCTCTCCCTGTACTGCCTAG |
| *Macaca fascicularis* | Cynomolgus monkey | ATGTACCCTTACGATGTACCTGACTACGCGGAGGCCCATAATGCGTC<br>TGCTCCTTTTAATTTCACATTGCCCCCAAATTTCGGGAAACGGCCGAC<br>GGATCTTGCCTTGTCCATTATTCTTGTATTCATGCTGTTCTTCGTCATG<br>TTGTCCTTGGGCTGTACAATGGAGTTCTCTAAAATCAAAGCCCATCTG<br>TGGAAGCCAAAAGGGCTCGCCATCGCGCTGGTGGCGCAGTACGGTAT<br>TATGCCTCTGACAGCGTTTGTGCTGGGAAAAGTCTTTCAACTCAACAA<br>TATAGAAGCTCTGGCTATTCTCGTGTGCGGGTGCAGTCCTGGGGGCA<br>ATTTGAGCAATGTGTTCTCCCTCGCGATGAAAGGAGACATGAACCTC<br>TCTATAGTCATGACTACCTGTTCCACCTTTTGTGCCCTGGGGATGATG<br>CCGCTCCTCCTGTACCTTTACACTCGGGGAATTTATGATGGCGACCTG<br>AAAGACAAAGTGCCCTATGGACGCATAATACTCTCCCTCGTGCCTGT<br>GCTTATCCCATGTACTATCGGTATAGTTTTGAAATCCAAGCGACCTCA<br>GTATATGAGGTATGTAATAAAGGGAGGCATGATTATTATTCTTCTCTG<br>CAGCGTCGCCGTAACTGTTTTGTCAGCTATAAATGTTGGCAAGTCCAT<br>AATGTTCGCAATGACGCCACTGCTGATTGCCACTAGTTCACTCATGCC<br>ATTCATCGGATTCCTTTTGGGGTATGTGTTGTCCGCCCTGTTTTGCCTT<br>AACGGTCGATGCAGACGCACTGTCAGCATGGAGACAGGCTGTCAGA<br>ACGTTCAACTCTGTAGTACCATACTGAATGTAGCATTCCCCCCCGAAG<br>TAATAGGGCCTCTTTTCTTTTTTCCTCTCCTTTACATGATCTTCCAACT<br>GGGGGAGGGACTCTTGCTCATAGCCATGTTTCGCTGCTACGAAAAGT<br>TTAAGACACCAAAGGACAAGACTAAAATGATATACACCGCCGCGACT<br>ACAGAGGAGACTATACCGGGGGCTCTCGGGAACGGTACTTACAAGG<br>GAGAAGATTGCTCACCGTGTACCGCGTAA |

| | | |
|---|---|---|
| *Saimiri sciureus* | Squirrel monkey | ATGTACCCTTACGATGTACCTGACTACGCGGATGCCCACAACATGAG<br>TGCGACGTTCAACTTCACATTGCCGCCAAACTTTGGAAAACGACCAA<br>CAGATCTCGCGCTGTCCATCATCCTCGTGTTCATGCTCTTCTTTATCAT<br>GTTGAGTTTGGGATGTACTATGGAATTCAGTAAGATTAAAGCTCATTT<br>TTGGAAGCCCAAAGGACTTGCAATTGCACTTGTAGCCCAATACGGTA<br>TTATGCCTCTTACGGCTTTCGTTCTGGGCAAAGTATTCCAGCTTAATA<br>AAATTGAAGCGTTGGCGATTTTGGTTTGCGGCTGTAGTCCGGGCGGT<br>AATCTTTCTAACGTGTTCTCATTGGCTATGAAAGGGGATATGAACCTC<br>TCAATCGTAATGACAACATGCAGTACGTTTTGCGCTCTCGGCATGATG<br>CCCCTTCTTCTCTATATATATTCTCGCGGCATTTATGATGGTGACTTGA<br>AGGATAAGGTTCCGTACGGAGGAATAATGATTTCTCTGATCCTGGTG<br>CTGATTCCATGTACCATAGGAATAGTTCTCAAATCTAAAAGACCCCA<br>ATATGTTCGGTATGTAGTGAAAGGGGGAATGATTATCATTCTGCTCTG<br>CAGTGTCACGGTCATCGTCTTGTCCGCTATTAATGTCGGAAAGTCTAT<br>ATTGTTCGCGATGACGCCTCTGCTTGTAACGACATCTTCCCTCATGCC<br>TTTCATTGGATTTCTGTTGGGTTACGTACTGTCCGCACTGTTCTGCCTC<br>AATGGTCGGTGTCGGCGCACTGTTTCCATGGAGACCGGCTGCCAAAA<br>TATACAACTCTGTTCAACCATTCTTAATGTTGCATTTCCCCCAGAAGT<br>TATAGGTCCACTCTTTTTTTTCCCTCTTCTTTACATGATTTTTCAGCTT<br>GGAGAGGGTCTCCTTCTCATAGCGATGTTTAGGTGTTACGAGAAATTT<br>AAGACGCCTAAGGACAAGACCAAGATTATCTACACTGCAGCTACTAG<br>CGAGGAAACAACTCCAGGAGCAGTTGGCAATGGCACCTACAAGGGC<br>AAAGAGTGTTCCCCATGCAAGGCTTAG |
| *Aotus nancymae* | Nancy Ma`s night monkey | ATGTACCCTTACGATGTACCTGACTACGCGGAGGCGCATAACGTCAG<br>TGCGGCTTTCAACTTTAGTTTGCCTCCAAACTTCGGCAAACGCCCGAC<br>TGATCTCGCGCTCAGTATAATACTGGTGTTTATGCTGTTTTTTATTATG<br>CTGAGCCTTGGCTGTACAATGGAGTTTTCAAAGATCAAAGCGCACTT<br>TTGGAAGCCTAAGGGCCTGGCGATCGCTCTGGTGGCGCAGTACGGGA<br>TAATGCCACTTACTGCGTTTGTTCTGGGTAAAGTTTTCCGCCTTAACA<br>AGATTGAAGCCCTCGCTATCCTCGTATGCGGTTGTTCACCCGGGGGTA<br>ACCTGTCCAACGTCTTCTCACTTGCTATGAAGGGAGACATGAATCTCT<br>CTATAGTAATGACAACATGCTCCACTTTCTGTGCTCTGGGGATGATGC<br>CCCTTCTCCTGTACATATATAGCCGGGGTATTTATGACGGTGACCTCA<br>AAGACAAGGTACCTTATGGGAGTATAATGCTCAGCCTGATTCTTGTTC<br>TTATCCCCTGCACCATTGGTATAGTATTGAAGTCCAAACGACCCCAAT<br>ACGTTCGCTATGTCGTGAAAGGAGGAATGATTATTATCCTTCTTTGCA<br>GCGTAACTGTGATAGTACTCTCCGCTATTAATGTTGGTAAGAGCATCC<br>TTTTTGCGATGACTCCTCTCTTGGTTACGACGTCTAGCCTCATGCCCTT<br>TATAGGATTTCTGCTGGGCTACGTGTTGAGCGCACTCTTTTGCCTGAA<br>CGGACGCTGTAGAAGGACTGTTTCTATGGAGACGGGTTGTCAGAATG<br>TGCAACTTTGCAGTACCATACTTAATCTCGCTTTTCCCCCAGAGGTTA<br>TTGGTCCCTTGTTTTTTTTTCCACTGTTGTACATGATATTCCAACTCGG<br>AGAGGGGTTGCTGCTGATCGCTACATTTAGGTGTTATGAAAAGTTTA<br>AAACTCCTAAGGATAAGACCAAGATAATCTACACAGCAGCAACCACT<br>GAAGAGACTACCCCCGGTGCAGTCGGGAATGGAACGTACAAGCGCA<br>AGGAGTGTTCTCCGTGTCGGGCATGA |
| *Felis catus* | Cat | ATGTACCCTTACGATGTACCTGACTACGCGGAACCACACAATGTGAC<br>GGCGACGCTCAACTTTACCCTCCCTCCGAACTTCGGGAAGCGCCCTA<br>CCGACAAAGCTCTGTCAGTTATACTTGTTTTCCTGCTCCTCATTATTAT<br>GCTGAGTCTGGGTTGCACCATGGAGTTTAGCAAAATACGAGCACACT<br>TTTGGAAACCAAAAGGACTTGCCATTGCCCTCATTGCACAATACGGC<br>ATCATGCCGTTGACAGCATTCGCACTGGGCAAGGTGTTTCAGCTGAA<br>CAACATTGAGGCGTTGGCCATTCTTGTGTGTGGTTGTAGTCCAGGTGG |

| | | |
|---|---|---|
| | | CACTCTGTCCAACATATTCTCACTGGCGATGAAGGGTGATATGAATCT<br>CAGCATTGTCATGACAACCTGTTCCACTTTCTGCGCGCTTGGGATGAT<br>GCCTCTGCTCTTGTATGTTTACAGTCGCGGAATCTACTCAGGTGACCT<br>CAAGGACAAAGTTCCTTATGGTGGGATTGTAATTAGTCTTATACTCGT<br>GCTCATCCCTTGTGCTACGGGAATTTTCCTCAATGCCAAGAGACCACA<br>ATACGTGCCATACGTAAAGAAAGGTGGAACGATCACGATGCTCCTGT<br>TGAGTGTAGCCATTATTGCTCTTAGCGTAATCAACGTGGGCAAATCTA<br>TTATGTTTGTTATGACACCCCACCTGCTGGCAACCTCTAGTTTGATGC<br>CGTTCATAGGCTTCCTTCTGGGATATATCTTGTCCGCACTTTTTCGCTT<br>GAACGGGCGGTGCCGCCGGACAGTATCTATGGAAACGGGCTGCCAG<br>AACGTGCAGCTTTGCTCTACCATTCTGAACGTGACTTTTCCGCCACAG<br>GTGATCGGGCCCCTCTTCTTTTTCCCACTCCTCTATATGATTTTTCAGT<br>TGGGAGAAGGGGTTCTGCTGATCTCCATATTCAGGTGTTATGAAAAA<br>ATGAAACCTAGTAAAGACAAAACCAAGATGACCTACACCGCCGCGC<br>CAACTGAAGATACAATACCGGGGGCCCTTGGGAATGGGACGCATAA<br>AGGTGAAGAGTGTTCACCCTGTACTGCTTAG |
| *Panthera tigris* | Tiger | ATGTACCCTTACGATGTACCTGACTACGCGGAACCACACAATGTTAC<br>CGCAACCCTTAACTTTACCTTGCCGCCTAATTTGGTAAGAGGCCTAC<br>GGACAAGGCCCTGAGTGTGATTCTCGTGTTCCTCCTTCTTATCATCAT<br>GCTCAGTCTGGGATGCACTATGGAGTTTAGTAAAATCAAAGCTCACT<br>TTTGGAAACCTAAGGGATTGGCTATTGCATTGATCGCACAATATGGC<br>ATCATGCCACTCACCGCCTTTGCATTGGGAAAGGTTTTCAGCTCAAC<br>AATATCGAAGCTCTTGCAATCTTGGTATGTGGGTGTTCTCCTGGAGGC<br>ACGCTTTCAAACATTTTTTCCTTGGCAATGAAAGGGGACATGAACCT<br>GTCCATTGTTATGACAACATGCTCTACGTTTTGTGCTCTCGGTATGAT<br>GCCGCTTTTGCTTTATGTGTATTCACGCGGTATATATGCGGGGGACTT<br>GAAGGATAAAGTCCCGTATGGTGGGATAGTGATCTCTCTCATACTGG<br>TTCTGATACCATGCGCTACAGGGATCTTTCTTAATGCTAAGAGGCCAC<br>AGTATGTACCCTATGTCAAAAAGGGTGGCACCATCACAATGCTCCTC<br>CTTAGTGTGGCGATCATAGCGCTCAGTGTAATTAATGTGGGAAAGAG<br>CATTATGTTTGTAATGACTCCCCACCTGCTCGCAACCTCCTCCCTGAT<br>GCCCTTCATAGGGTTCCTCTTGGGATATATTCTGAGTGCCCTGTTCCG<br>CCTTAACGGGCGATGCCGGAGAACCGTTAGTATGGAAACCGGGTGCC<br>AAAATGTCCAATTGTGTTCAACGATCCTTAACGTGACTTTTCCCCCCC<br>AAGTGATCGGACCCTTGTTCTTTTTTCCTTTGCTGTACATGATATTCCA<br>GTTGGGGGAGGGGGTTTTGCTGATCTCCATTTTCCATTGCTACGAAAA<br>GATAAAACCTAGCAAGGACAAGACAAAAATGACATATACTGCAGCT<br>CCGACCGAAGACACCATACCAGGAGCACTTGGCAATGGCACACATA<br>AAGGCGAAGAATGCAGCCCTTGCACGGCATAA |
| *Leopardus geoffroyi* | Geoffroyi's cat | ATGTACCCTTACGATGTACCTGACTACGCGGAGCCCCTCAACGTAAC<br>AGCGACCCTGAATTTTACACTTCCACCAAACTTTGGCAAGAGACCTA<br>CAGATAAGGCACTCTCCGTTATTCTGGTATTCTTGTTGTTGATAATTA<br>TGCTGTCACTGGGATGTACAATGGAATTTTCCAAGATCAAGGCTCAC<br>TTCTGGAAACCAAAGGGTTTGGCCATCGCGCTTATAGCCCAATACGG<br>TATAATGCCGCTGACTGCGTTCGCTCTGGGTAAAGTATTCCAGCTGAA<br>CAATATTGAGGCTCTTGCCATTCTTGTTTGCGGTTGTAGCCCTGGCGG<br>CACACTCTCCAATGTGTTCTCCCTGGCCATGAAGGGCGATATGAACCT<br>TTCTATAGTCATGACAACTTGTAGTACCTTCTGTGCCCTTGGGATGAT<br>GCCGCTTCTCTTGTACGTTTATTCCAGAGGCATTTATGCGGGCGACCT<br>TAAGGACAAAGTCCCGTACGGGTCAATTGTCATTAGCCTTATCCTCGT<br>CCTTATTCCTTGCGCTACTGGTATCTTCCTGAATGCCAAGAGACCCCA<br>ATACGTACCGTATGTTAAAAAAGGGGGCACGATAACTATGTTGCTCC<br>TTAGTGTGGCGATCATAGCCCTCTCAGTGATAAACGTTGGGAAGTCC |

| | | |
|---|---|---|
| | | ATTATGTTTGTGATGACACCTCATCTTTTGGCAACGTCATCACTGATG<br>CCTTTTATAGGATTCCTCCTTGGCTATATTCTGTCAGCGTTGTTCCGAT<br>TGAACGGGAGGTGCCGGCGGACTGTCTCTATGGAAACCGGCTGTCAG<br>AACGTACAACTTTGCAGCACCATCTTGAATGTGACTTTCCCGCCGCAG<br>GTCATAGGCCCCCTGTTCTTTTTCCTCTGCTTTATATGATCTTTCAAT<br>TGGGCGAAGGTGTTCTGCTTATTAGCATTTTTCGATGCTATGAAAAAA<br>TAAAACCCTCTAAGGATAAAACGAAGATCACTTATACAGCAGCGCCA<br>ACGGAGGACACGATCCCGGGGGCTCTTGGGAACGGGACACATAAGG<br>GAGAGGAGTGTTCACCGTGCACGGCGTGA |
| *Canis lupus familiaris* | Dog | ATGGACGCGCCAAACATCACTGCACCGCTTAACTTCACTCTGCCTCCC<br>AACTTCGGTAAACGCCCCACAGATAAGGCTCTTTCCATCATTCTTGTA<br>TTCCTGTTGCTTATAATTATGCTTAGCCTCGGGTGTACGATGGAATTT<br>AGCAAAATTAAGGCGCATTTTTGGAAACCCAAGGGTCTGGTCATTGC<br>TCTGATTGCCCAATATGGAATTATGCCCCTGACGGCCTTCACGCTTGG<br>CAAGGTGTTCAGGCTGAATAATATCGAGGCACTGGCTATCTTGGTTT<br>GTGGCTGCTCTCCCGGCGGAACCTTGTCTAACGTGTTTAGTTTGGCTA<br>TGAAGGGCGACATGAACCTGTCCATTGTTATGACCACGTGCAGTACG<br>TTTTTTGCTCTGGGTATGATGCCCCTGCTGCTCTATATATATAGCAAC<br>GGTATCTACGACGGTGACTTGAAGGACAAGGTGCCATACAAGGGAAT<br>TGTTTCCTCTCTCGTTCTCGTTTTGATACCCTGCACGATAGGCATTTTT<br>TTGAAAGCCAAACGCCCACAGTACGTACGATACATAAAGAAGGGCG<br>GTATGATAATCATGCTGCTTTTGTCTGTTGCGATCACGGCGCTTTCTG<br>TAATTAACGTAGGAAAGTCAATACGATTCGTGATGACTCCTCATCTGT<br>TGGCGACCAGTAGCCTGATGCCTTTTATCGGGTTTCTTCTCGGCTACA<br>TCTTGAGCGCCCTTTTCCGCCTTGACGGTAGGTGCAGTCGGACGGTGA<br>GTATGGAAACCGGTTGTCAGAATGTTCAACTGTGCTCCACGATACTC<br>AACGTAACCTTCCCCCCTGAGGTTATCGGACCGCTGTTTTTTTCCCC<br>CTCTTGTATATGATATTCCAACTCGGCGAGGGGGTATTTTTGATTTCC<br>ATATTTAGATGCTACGAAAAAATTAAGCCGAGTAAAGATAAGACCAA<br>AATGATATATACGGCGGCAGCAACCGAGGAAATCACGCCGGGGGCA<br>CTCGGGAATGGAACGCATAAGGGTGAAGAATGCAGCCCGTGCACCG<br>CTGCGCCCTCACCTAGTGGTCTTGACTCCGGCGAAAAGGCTATTCAGT<br>GTGATCAGCTCGAAAAGGCGAAGGACAAGAGAAACACCAAAGAGGA<br>AAGTTTCTCTAGCATAGGCTCCTCAAATTATCAGAATTAG |
| *Rhinolopus ferrumequinum* | Horseshoe bat | ATGTACCCTTACGATGTACCTGACTACGCGGAGGCGCACAATGGCTC<br>CGCTCCATTGAACTTTACTCTGCCGCCCAATTTCGGTAAAAGACCGAC<br>AGACCTCGCTCTCAGCGTTATCCTCGTATCCATGCTGCTCATCATGAT<br>GCTCAGTCTTGGCTGCACGATGGAGTACTCAAAAATAAAAGCCCATT<br>TTTGGAAACCAAAGGGATTGGCCATAGCTCTGGTGGCCCAATACGGG<br>ATAATGCCTTTGACGGCTTTTCTGATTGGTAAGATATTCCGCCTCAAT<br>AATATAGAGGCTTTGGCTATCTTGATTTGTGGCTGTAGCCCAGGAGG<br>AAACCTGAGCAACGTCTTTTCATTGGCGTTGAAGGGCGATATGAATC<br>TGAGTATTGTGATGACAACTTGTTCTACTTTTTTTGCCCTGGGCATGA<br>TGCCACTTTTGCTGTATATCTACTCTAGAGGGATTTATGACGGTGATC<br>TTAAAGAAAAAGTTCCATATGGCGGTATCGTTTTGTCATTGGTTCTCA<br>TATTGATACCATGCACCATAGGGATATTCTTGAACGCAAAGCGACCT<br>CAATACGTAAGGTACGTGACAAAAGGGGGTATGATAATCACTTTGCT<br>TTTGTCCGTCGCTGTCATTGCTCTTTCCGCCATAAACGTGGGAAAGTC<br>AATCATGTTTGTCATGACACTTCGCCTCTGGGCGACATCCTCATTGAT<br>GCCATTTATAGGATTTTTGCTGGGATATATCCTGAGCGTCCTTTTTAG<br>GCTCAACGGGCAATGTTCCAGAACAGTCTCAATGGAAACTGGCTGCC<br>AGAATGTTCAACTCTGCAGCACGATTCTGAACGTCACTTTCCCGCCTG<br>AAGTAATTGGTCCTCTGTTTTTCTTCCCATTGCTTTATATGATTTGTCA |

| | | |
|---|---|---|
| | | ATTGGGGGAGGGGCTTCTCCTTATTGCCATCTTTCGGTGTTATGAGAA<br>AATTAAGCCAAGCAAGGACAAGACCAAGATTATATATAAGGCGGCA<br>ACTACAGCAGAAACTACAGTTCCGTCTCCACTCGGCAACGGTACGCA<br>CAAAGAAGGATTTCCACGAACGGCTTAA |
| *Pteropus alecto* | Black fruit bat | ATGTACCCTTACGATGTACCTGACTACGCGGAAGCCCATAATGCGTC<br>TGCTCCTCTTAACTTCACATTGCCTCCCAATTTTGGGAAGCGCCCCAC<br>TGACCTCGCCGTGTCTATTATCCAAGTGTTTATGCTTCTCATTATGAT<br>GTTGTCCCTCGGTTGCACCATGGAGTTTTCCAAGATAAAAGCACACTT<br>TTGGAAGCCACGGCGGTTGGCCATCGCTCTGATGGCCCGGTATGGCA<br>TTATGCCTCTTACGGCCTTCGTCCTGGGCAAAGTGTTCAGGCTGAATA<br>ACATTGAAGCACTCGCGATACTTATATGCGGATGTAGCCCCGGCGGT<br>AACCTGTCTAATTTGTTTTCCCTCGCCATAAAAGGAGATATGAATTTG<br>AGAAAGGATTGGGGTATAATGATAACCACTTGTTCAACGTTTCTGGC<br>GCTTGGCATGATGCCGCTCCTGTTGCACGTGTATTCTAGGGGGATCCA<br>TGACGGGGATCTGAAAGATAAAGTCCCTTATCGGGAGATAGTTCTCT<br>CTTTGGTTCTCATTTTGATACCGTGTACGATAGGCATATTCCTCAACG<br>CAAAGAGGCCACAGTACGTGTGCTACATCATTAAAGGCGGTAGAATA<br>ATTACTCTGCTGTTGAGCGTTGCGATCACGGCACTGTCAGCTATCAAC<br>GTGGGAAAATCAATTCTGTTCGTCATGACACCCAGGCTGTGGGCGAC<br>CAGTTCCTTGATGCCCTTCATTGGGTTTCTCCTCGGATACATCCTCAG<br>TGCTTTGTTTTATCTTTCTGGCGACTGCCGCCGGATCGTATCTATGGA<br>AACGGGGTGCCAAAATATACAACTTTGTTCCACGATCCTTAACGTGA<br>CACTGCCTCCGGAAGTTATCGGCCCTCTGTTCTTTTTCCACTCCTTTA<br>CATGATTTGCCAAGTAGCGGAGAGGTTTCTTTTGGTTGCAATGTTTAG<br>GTGTTACGAGAAAATTAACCCCTCAAAGGATAAGACTATCTACACCG<br>CTGCTACAACAGAGGAAATCATTCCCGCCGCACTTGAAAATAAAGAA<br>GAATGCTCTCCGGGAACAACCTAG |
| *Sus scrofa* | Pig | ATGTACCCTTACGATGTACCTGACTACGCGGAGGCACTCAATGAATC<br>AGCCCCAATTAACTTCACATTGCCTCATAACTTTGGGAAACGACCAA<br>CAGATTTGGCACTCAGTGTGATACTTGTTTTTATGCTCTTGATAATAA<br>TGCTCAGCCTTGGATGTACGATGGAATTTGGTAGAATCCGGGCACAT<br>TTCAGAAAACCGAAAGGTCTGGCAATCGCATTGGTAGCTCAGTACGG<br>TATCATGCCGCTGACAGCGTTCGCTCTGGGCAAGTTGTTTCGATTGAA<br>CAATGTGGAGGCCCTTGCCATACTTATCTGCGGTTGTAGCCCCGGCG<br>GGAATCTGAGTAATATTTTCGCTCTCGCGATGAAAGGTGATATGAAT<br>CTGAGCATCATGATGACCACCTGCAGTACTTTCCTGGCCCTGGGGAT<br>GATGCCGCTCCTGCTTTATCTGTATTCCCGAGGGATCTACGACGGCAC<br>CTTGAAAGATAAAGTACCCTATGGGTCTATAGTGATATCCCTCATCCT<br>GATTCTTATACCGTGCACAATAGGCATAATCTTGAACACAAAACGGC<br>CACAATATGTCCGGTATGTAATTAAAGGCGGTACTATACTTCTGATTC<br>TTTGTGCAATAGCAGTCACAGTACTGTCAGTCTTGAATGTTGGGAAGT<br>CAATTCTCTTCGTCATGACTCCTCACCTCGTCGCGACAAGTAGCCTGA<br>TGCCATTTACTGGGTTCCTGCTCGGATACCTGCTTAGCGCTCTGTTTC<br>GGTTGAATGCTCGGTGTTCCCGAACCGTGTGCATGGAGACAGGATGC<br>CAAAACGTACAACTGTGCTCAACAATTCTGAACGTGACGTTTCCTCCC<br>GAGGTCATCGGTCCCTTGTTTTTCTTCCCACTCCTCTATATGCTGTTTC<br>AACTTGGAGAAGGACTGCTTTTCATAGCTATCTTCCGCTGCTATGAGA<br>AAACCAAATTGAGCAAGGACAAAATGAAAACAATCTCTGCTGCAGA<br>CAGTACCGAGGAGACAATTCCGACCGCTCTTGGCAATGGGACACACA<br>AGGGTGAGGAATGTCCCCCAACTCAACCGTCCGTGGTTTAG |
| *Bos taurus* | Cow | ATGTACCCTTACGATGTACCTGACTACGCGGAGGCGTTCAATGAGTC<br>AAGTCCCTTCAATTTCTCACTTCCACATAACTTTGGGAAGCGGCCGAC<br>AGACAGGGCCTTGAGTGTCATTCTCGTCATCATGCTTCTGACAATTAT |

| | | |
|---|---|---|
| | | GCTCAGTCTCGGGTGTACCATGGAGTTTTCCAAAATTAAAGCTCACTT<br>CTGGAGACCTAAAGGATTGGCGGTCGCACTTGTCGCCCAATTTGGAA<br>TCATGCCCCTGACTGCCTTCGGACTGGGCAAGTTTTTCAACTTAATA<br>ATGTAGAGGCTCTTGCCATTCTGATATGTGGTTGTAGTCCGGGCGGA<br>AATCTTTCCAATGTTTTCGCGCTTGCCATAAAGGGTGACATGAACCTC<br>TCAATTGTCATGACTACTTGTAGCACTTTCTTCGCACTGGGAATGATG<br>CCGTTGCTGTTGTATCTCTATAGTAGAGGCATTTACGATGGGTCACTG<br>AAAGACAAGGTTCCTTATGGTGGCATCATGATCTCTTTGATCCTTATA<br>CTCATACCGTGTACTATTGGTATTATTTTGAAGTCTAAGCGGCCTCAA<br>TACGTCCGATATGTGACGAAAGGTGGGATCATTCTCCTGTTGCTGTGT<br>TCCGTTGCAGTAGTTGTGCTCAGCGCAATAAACGTTGGTAAAAGTAT<br>ATTGTTCGTGATGACACCCCATTTGCTTGCCACGTCTTCCTTGATGCC<br>GTTCATTGGCTTTCTTTTGGGATACCTTCTTAGCGCTTTGTTCTGCCTG<br>AACGGGCGCTGTAAGCGAACAGTCAGCATGGAAACTGGCTGTCAAA<br>ACATCCAGCTCTGCTCTACAATACTGAATGTAACATTTCCGCCCGAAG<br>TTATAGGTCCTCTTTTTTTCTTTCCTCTTCTTTATATGATATTTCAGGTA<br>GGCGAGGGCCTGCTTTTGGTTGCGATTTTCAGATGCTATGAAAAGTTT<br>AAGACCCCAAGGGATGATACCAAAATGACGTACAAAGTTGACGCTA<br>CAGAAGAGACATTGCCCACTGCGCTGGGCAATGGCACATACAAGGGT<br>GAGGAGTGTAATATTGGTTTTACAAGCGGCTTCCCTAAGGAGCTGGA<br>CTCCGGCCAAAAAGCTACAGAACAACTCAACATGGCCAACTAG |
| *Tursiops truncates* | Dolphin | ATGTACCCTTACGATGTACCTGACTACGCGGGCTTGCTCGGCCTCTTG<br>GCTAAGGAACCTGGTCAAAAGAGCATTTCTCCTGCGCGGGCACCAAG<br>TGAATTGCCGTTTGGGGAGAGAGCACCAAGCTTGCCCGAGTCCCCCA<br>GTGCTTCAGCGGCAGGGTCAAGCCTTGAACTCGGACAAGGCTCCTGG<br>ACACAACTGCTGCTTGAGGCGAGTAATAGCCCTGCGGCGACACTCAG<br>TCACACACCTCTCTCACGCAAGGAGCGACGGATGGAGGCTCTTAACG<br>AATCCGCGCAGTTCAATTTCTCCCTTCCTCATAATTTCGGAAAAAGGC<br>CTACCGACTGGGCATTGAGTGTCATTTTGGTAATCATCCTGCTTACTA<br>TAATGTTGTCTCTTGGCTGCACGATGGAATTTAGCAAAATAAAAGTG<br>CACTTCTGGAAACCGAAAGGGCTTGCTATCGCTTTGGTGGCCCAATA<br>TGGGATCATGCCATTGACCGCTTTCGCCTTGGGCAAAGTGTTTCAACT<br>TAACAATGTAGAGGCTCTGGCAATCCTGATTTGCGGGTGTTCCCCGG<br>GTGGCAATTTGTCAAATGTATTCACGCTTGCTATGAAGGGGGACATG<br>AATCTTTCTATTGTCATGACGACATGTTCTACCTTCTTCGCCTTGGGA<br>ATGATGCCTTTGCTGCTCTATATCTACTCCAGAGGCATATACGACGGA<br>ACCTTGAAAGATAAGGTTCCTTATGGCAACATAATGATTTCTCTCATA<br>CTCATACTGATTCCTTGCACTATCGGTATTATTCTGAAGAGTAAGAGG<br>CCCCAATACGTCCGGTATGTGACCAAGGGGGGGATGATTTTGTTGTT<br>GCTTTGCTCAGTGGCCGTGGTAGTTCTGTCCGTACTTAACGTCGGGAA<br>GAGTATTCTGTTTGTGATGACACCACATCTTCTGGCGACGAGTAGTCT<br>CATGCCGTTCATAGGCTTCCTGCTGGGTTATGTGCTGTCAGCTCTTTT<br>CCGACTCAATGGGCGGTGTAGGCGCACGGTGTCCATGGAGACCGGTT<br>GCCAAAATATACAGCTGTGTAGCACTATTCTGAACGTTACGTTCCCGC<br>CCGAGGTAACGGGGCCGCTGTTTTTCTTTCCACTGCTGTACATGATCT<br>TTCAACTGGGGGAGGGGTTGCTGCTCATCGCTATCTTCAGGTGTTATG<br>AGAAGATCAAGCCCAGCAAGCACAAGACGAAAATGATTTATAAAGC<br>TGATACGACAGAAGAAACGATACCAACAGCGTTGGGCAATGGTACA<br>CACCGCGGTGAAGCGTGCCCACCGTGTACAGCGTGA |
| *Marmota monax* | Woodchuck | ATGTACCCTTACGATGTACCTGACTACGCGGAAGTTTACAATGTGAG<br>CGTCCCGTTCAATTTTAGTCTCCCGCCGAACTTTGGTAAGAGGCCGAC<br>CGATCTGGCGCTCTCTATTATATTGGTGTTCATGCTCCTGATCATAAT<br>GCTGTCACTCGGCTGCACGATGGAGTTCTCCAAAATCAAAGCACATC |

| | | |
|---|---|---|
| | | TTTTGAAACCAAAGGGTCTTGCAATAGCCATGGTAGCACAGTACGGG<br>ATAATGCCGTTGACAGCCTTTGTGCTGGGGAAGGTGTTCCAATTGAA<br>TAACATTGAAGCACTGGCAATACTGATTTGCGGGTGTTCTCCCGGTG<br>GCAACTTGAGCAATATCTTTTCCTTGGCGGTAAAGGGAGACATGAAC<br>CTGTCCATAGTCATGACTACATGTAGCACTTTTTCGCCCTCGGTATG<br>ATGCCGCTCCTGCTCTATATATATTCCAAGGGCATATACGACGGAAG<br>TCTTGAAGATAAAGTTCCATATAAAGGTATCATGATTTCCTTGGTCAT<br>GGTACTCATCCCTTGTACTATAGGAATTATCCTGAAATCTAAGAGACC<br>CAAGTATGTCCCCTATATCATGAAAGGTGGTATGATCATAACACTCTT<br>GTTTAGTGTAGCGGTAACGGCCCTCAGTATCATCAACACTGGCAAGT<br>CTATAATGTACGCAATGACCCCTCATCTCTTGACTGTGTCCTCATTGA<br>TGCCGTTCATAGGGTTCCTGCTGGGATATATACTCTCAGCCTTGTTCT<br>GTTTGAAAGCTCGATGCCGCAGGACGATTAGCATGGAGACAGGTTTC<br>CAGAATATCCAGCTCTGTTCTACCATATTGAATGTAACATTTCCACCA<br>GAAGTTATAGGACCTCTCTTTTTTTTCCCGTTGCTGTACATGATATTCC<br>AACTTGGCGAGGGCCTGCTTTTCATCATAGTTTTCCGGTGTTACAAGA<br>AATTCAACCCACCAAAAGAAAAAACCAAGATGATTTATACAGCGGC<br>GACGAAAGAAGCTATTTCTCGCGCGCAAGAAAACGGTGTACACAAG<br>GGAGAAGAATGCTCACCTTGCACTGCCTGA |
| *Incidomys tridecemlineatus* | Ground squirrel | ATGTACCCTTACGATGTACCTGACTACGCGGAGGTTCATAATGTTAGC<br>GTACCTTTTAATTTTTCTCTGCCGCCTAATTTCGGGAAGCGGCCAACG<br>GACCTGGCTTTGTCTATTATACTTGTATTGATGCTTCTGGTGATTATGC<br>TCTCTCTTGGGTGCACTATGGAGTTTAGCAAGATTAAAGCACATTTGT<br>TGAAGCCTAAGGGCCTTGCAATTGCTATGGTGGCCCAGTACGGTATC<br>ATGCCACTCACTGCATTTGTACTTGGCAAGGTATTCCAACTCAACAAC<br>ATAGAGGCCCTTGCGATACTGATTTGTGGATGCTCCCCAGGCGGGAA<br>CCTCTCCAATATATTCTCCCTTGCAGTTAAGGGAGACATGAACCTCAG<br>CATCGTAATGACCACCTGCTCCACTTTCGTGGCGCTGGGTATGATGCC<br>GCTCTTGCTTTATATTTATAGTAAAGGGATTTACGACGGGAGCCTCGA<br>GGACAAGGTACCATATAAAGGAATCATAATCTCTCTGATTATGGTAC<br>TGATCCCTTGTACAATTGGGATAATACTTAAATCTAAACGACCGAAG<br>TACGTTCCTTACATAATGAAAGGCGGTATGATAATAAGCCTGTTGTTC<br>AGTGTCGCCGTGACCGCTCTTAGCATTATTAACATAGGAAAGTCTATT<br>ATGTACGCCATGACCCCTCACTTGTTGACAGTAAGCTCACTTATGCCA<br>TTCATTGGGTTCTTGTTGGGTTATTTGCTCTCAGCTCTGTTTTGTTTGA<br>AAGCTCGATGCAGAAGAACGATTAGCATGGAAACGGGGTTTCAGAA<br>TATACAATTGTGTTCTACAATACTCAACGTGACGTTTCCACCGGAGGT<br>AATTGGCCCTCTCTTTTTCTTCCCCTTGCTGTACATGATCTTCCAGCTT<br>GGCGAGGGACTCCTGTTCATCATTGTTTTTCGCTGTTACAAGAAATTC<br>AACCCACCCAAGGAAAAGACTAAGCTCATTTACACAGCCGCAACCAA<br>GGAAGCCATATCTCGGGCGCAAGAAAATGGTGTTCACAAAGGGGAA<br>GAGTGCACGCCGTGCACGGCGTGA |
| *Mus musculus* | Mouse | ATGTACCCTTACGATGTACCTGACTACGCGGAAGCGCATAATGTCTC<br>CGCTCCATTTAATTTTAGCTTGCCGCCCGGATTTGGGCATCGCGCAAC<br>TGATACCGCGCTGAGCGTTATCCTCGTAGTGATGCTCCTTCTCATAAT<br>GCTGAGCCTCGGTTGTACAATGGAATTCTCTAAGATTAAGGCTCACTT<br>CTGGAAGCCGAAGGGTGTGATCATAGCTATTGTGGCTCAATACGGCA<br>TAATGCCGCTCTCTGCTTTTCTTTTGGGGAAGGTGTTTCACCTTACATC<br>TATCGAGGCATTGGCAATTCTTATATGTGGGTGTTCCCCCGGAGGTAA<br>TCTCTCTAATCTCTTTACTCTTGCCATGAAGGGTGACATGAATCTCTC<br>TATCGTTATGACAACTTGTAGCTCCTTTACAGCACTTGGCATGATGCC<br>CTTGCTCCTCTATATCTATTCTAAGGGGATTTACGACGGAGACCTTAA<br>GGACAAGGTCCCATATAAGGGAATAATGCTCTCCTTGGTAATGGTGT |

| | | |
|---|---|---|
| | | TGATACCATGTGCAATTGGGATTTTCCTGAAAAGTAAGAGGCCACAT<br>TATGTCCCTTATGTGCTCAAAGCCGGCATGATTATTACTTTTAGCCTC<br>TCAGTTGCGGTTACGGTCCTGTCAGTCATTAACGTTGGCAATTCCATT<br>ATGTTTGTAATGACTCCCCATCTCTTGGCTACGTCATCACTGATGCCT<br>TTTACAGGCTTCCTCATGGGCTACATCTTGTCCGCTCTGTTCAGACTC<br>AATCCCTCTTGCCGACGAACGATTTCTATGGAGACAGGGTTCCAAAA<br>CGTACAATTGTGTTCCACGATCCTTAATGTTACGTTTCCGCCAGAGGT<br>CATTGGACCCTTGTTTTTTTTCCCCTCTTGTATATGATATTTCAGTTG<br>GCAGAGGGGCTGCTTTTCATCATAATCTTTCGCTGCTACCTTAAAATC<br>AAGCCACAGAAGGACCAGACTAAGATCACATACAAAGCGGCCGCCA<br>CGGAGGACGCTACTCCGGCCGCTCTGGAGAAGGGGACCCATAACGG<br>CAATAACCCGCCAACCCAACCCGGACTCTCTCCTAATGGGCTTAATA<br>GTGGCCAAATGGCGAATTAA |
| *Rattus novegicus* | Rat | ATGTACCCTTACGATGTACCTGACTACGCGGAGGTTCATAATGTCAG<br>CGCACCATTTAATTTTTCTCTCCCTCCAGGATTCGGCCATAGGGCCAC<br>TGATAAAGCCTTGTCAATAATCCTTGTTTTGATGTTGCTTCTTATTATG<br>CTCAGCCTGGGTTGCACAATGGAGTTTAGTAAGATTAAAGCCCATCT<br>CTGGAAACCGAAGGGGGTAATTGTTGCACTGGTGGCTCAGTTTGGAA<br>TAATGCCTCTTGCGGCGTTCCTCCTCGGCAAAATTTTTCATCTTAGCA<br>ACATAGAAGCACTGGCGATTCTGATTTGTGGGTGCAGTCCCGGTGGC<br>AACCTCTCTAACCTGTTCACCCTCGCGATGAAAGGTGATATGAATCTT<br>TCAATCGTTATGACAACCTGCAGTTCTTTCTCTGCCCTCGGTATGATG<br>CCTTTGTTGCTCTACGTCTACAGCAAAGGAATATATGACGGGGACCT<br>GAAAGATAAAGTACCATACAAAGGTATCATGATCAGTTTGGTGATAG<br>TACTGATCCCATGCACTATTGGGATTGTGCTCAAGAGCAAGCGACCT<br>CATTACGTCCCCTATATCCTCAAAGGTGGCATGATCATAACATTCCTC<br>TTGTCTGTAGCAGTGACGGCCTTGTCTGTAATCAATGTGGGAAACTCC<br>ATCATGTTCGTAATGACGCCCCATTTGTTGGCAACGAGTTCCCTTATG<br>CCGTTCTCTGGGTTTCTGATGGGCTATATATTGAGTGCTCTCTTTCAAT<br>TGAACCCGTCATGCAGGAGAACTATTTCCATGGAGACAGGCTTTCAG<br>AACATACAGCTTTGTAGTACGATCTTGAATGTAACATTCCCCCCTGAG<br>GTAATAGGCCCTCTCTTCTTTTTCCCGCTTCTCTACATGATCTTTCAAC<br>TGGCTGAAGGACTCTTGATAATAATTATATTCAGGTGTTATGAGAAG<br>ATAAAACCTCCTAAAGACCAAACGAAAATTACGTATAAAGCAGCGG<br>CAACCGAGGACGCGACACCGGCAGCCCTGGAGAAAGGAACCCATAA<br>TGGAAATATCCCTCCTTTGCAACCGGGTCCAAGTCCTAATGGTCTTAA<br>CAGTGGCCAGATGGCTAACTGA |
| *Ornithorhy cus anatinus* | Platypus | ATGTACCCTTACGATGTACCTGACTACGCGGAACTCCAAGGGCCGGC<br>CGACTCTAATGGCTCTTCTTCACTCAATTTTACGTTCCCCCCAAACTTC<br>GGAAGGCGGCCAAGTGATAAGGCACTGTCCATATCTCTGGTGGTGAT<br>GCTGTTCGTCACTATGATATCTCTCGGCTGCACCATGGAGTTCGCCAA<br>GATAAGGGCCCACCTGTTCAAGCCGAAGGGCGTGGCTATTGCCCTCG<br>TAGCGCAATATGGGGTCATGCCATTGACGGCCTTCACACTCGGAAGG<br>GTATTCCAATTGAATACCATAGAAAGTCTGGCAATTTTGATTTGCGGG<br>TGCTCCCCTGGCGGGTCCCTCAGTAACGTTTTCTCTCTTGCCATGAAA<br>GGTGACATGAACCTTTCAATTGTAATGACGACCTGCAGTACATTTAGT<br>GCGCTCGCCCTGATGCTGTTGTTGTTGTATCTCTACTCACTCGGCTTGT<br>ACGAGGGGGATATTCAAAATAAGGTACCATACGGTGGAATAATCCTT<br>TCACTCGTTCTCGTATTGATTCCATGTACTATTGGAATCTTCCTGAAA<br>TCAAAGAGACCGCAATACGTTCCGTATGTAATTAAGGTGGGGGCTAT<br>AATAACAGTCCTGATGCTTATAGCGGTTGTAGTGTTGTCTATTATCAA<br>CGTCAACCACAGTATACTGCTGATTATGACTGCACCCCTGCTGGCTAT<br>ATCTTCATTGATGCCGTTTACCGGTTTCTTCCTCGGGTACCTTCTCTCC |

| | | |
|---|---|---|
| | | GCACTGTTCCGACTGAACGGTCGGTGCCGAAGGACCGTCAGTATGGA GACTGGCTGCCAAAATGTCCAACTCTGCTCTACAATTCTTAATCTCAC CTTCCCCCTCGAGGTAATTGGTCCCTTGTTCTATTATTTGCTTTTGTAT ATTATATTCCAGGTTGGGGAGGGACTCATTTTGATTGCGCTCTTTCGC TGTTACGAGAAGATTCGGACTCCGCACGACGACCCAAAAATGATATA CAGAGCGGTAAGCGAGGCTGCATCTGAAGCGAAAACACGACCCAGA TCCAATGGTTCCCACGAAGGAGAGGAGAGGTCCATGAACTCCTTTCA TAGTCCGGCCGTTCCCACACTCCCTCTCGAACATTCCTCTCAAGGAGC GTCCAGCCAAGCCTAG |

**Supplementary Table 2. Primers used for mutagenesis of the residues 158 of NTCP**

| Species | Common name | Mutation introduced | Direction | Sequence (5′-3′) |
|---|---|---|---|---|
| *Homo sapiens* | Human | G158R | Forward | CCCTACAAACGTATCGTTATCTCACTCGT |
| | | | Reverse | GATAACGATACGTTTGTAGGGCACCTTGT |
| *Macaca fascicularis* | Cynomolgus monkey | R158G | Forward | CCCTATGGAGGCATAATACTCTCCCTCGT |
| | | | Reverse | GAGTATTATGCCTCCATAGGGCACTTTGT |
| *Aotus nancymae* | Nancy Ma`s night monkey | S158G | Forward | CCTTATGGGGGTATAATGCTCAGCCTGAT |
| | | | Reverse | GAGCATTATACCCCCATAAGGTACCTTGT |
| *Felis catus* | Cat | G158R | Forward | CCTTATGGTCGGATTGTAATTAGTCTTA |
| | | | Reverse | AATTACAATCCGACCATAAGGAACTTTGT |
| *Leopardus geoffroyi* | Geoffroyi's cat | S158G | Forward | CCGTACGGGGGAATTGTCATTAGCCTTAT |
| | | | Reverse | AATGACAATTCCCCCGTACGGGACTTTGT |
| *Pteropus alecto* | Black fruit bat | E158G | Forward | CCTTATCGGGGGATAGTTCTCTCTTTGGT |
| | | | Reverse | GAGAACTATCCCCCGATAAGGGACTTTAT |
| *Sus scrofa* | Pig | S158G | Forward | CCCTATGGGGGTATAGTGATATCCCTCAT |
| | | | Reverse | TATCACTATACCCCCATAGGGTACTTTAT |
| *Tursiops truncates* | Dolphin | N158G | Forward | CCTTATGGCGGCATAATGATTTCTCTCAT |
| | | | Reverse | AATCATTATGCCGCCATAAGGAACCTTAT |

**Supplementary Table 3. Estimated evolutionary divergence between mammalian NTCP sequences.**

| | Amino Acid | | | | | | | | | | | | | | | | | | | | |
|---|---|---|---|---|---|---|---|---|---|---|---|---|---|---|---|---|---|---|---|---|---|
| | | | 1 | 2 | 3 | 4 | 5 | 6 | 7 | 8 | 9 | 10 | 11 | 12 | 13 | 14 | 15 | 16 | 17 | 18 | 19 | 20 |
| Nucleotide | 1 | *Homo sapiens* | | 0.04 | 0.08 | 0.09 | 0.18 | 0.17 | 0.19 | 0.18 | 0.16 | 0.15 | 0.21 | 0.19 | 0.17 | 0.16 | 0.23 | 0.20 | 0.21 | 0.25 | 0.25 | 0.37 |
| | 2 | *Macaca fascicularis* | 0.03 | | 0.08 | 0.09 | 0.19 | 0.19 | 0.21 | 0.19 | 0.18 | 0.17 | 0.23 | 0.20 | 0.18 | 0.18 | 0.22 | 0.21 | 0.21 | 0.27 | 0.27 | 0.35 |
| | 3 | *Saimiri sciureus* | 0.06 | 0.07 | | 0.04 | 0.19 | 0.19 | 0.21 | 0.19 | 0.19 | 0.18 | 0.24 | 0.19 | 0.17 | 0.19 | 0.25 | 0.21 | 0.21 | 0.28 | 0.28 | 0.38 |
| | 4 | *Aotus nancymaae* | 0.06 | 0.07 | 0.03 | | 0.20 | 0.20 | 0.22 | 0.19 | 0.20 | 0.19 | 0.23 | 0.20 | 0.18 | 0.18 | 0.25 | 0.22 | 0.22 | 0.27 | 0.28 | 0.38 |
| | 5 | *Felis catus* | 0.14 | 0.15 | 0.16 | 0.16 | | 0.01 | 0.03 | 0.02 | 0.11 | 0.16 | 0.21 | 0.24 | 0.19 | 0.18 | 0.24 | 0.25 | 0.26 | 0.26 | 0.26 | 0.40 |
| | 6 | *Panthera tigris* | 0.14 | 0.15 | 0.17 | 0.16 | 0.01 | | 0.02 | 0.01 | 0.11 | 0.15 | 0.22 | 0.24 | 0.19 | 0.18 | 0.23 | 0.25 | 0.26 | 0.25 | 0.26 | 0.41 |
| | 7 | *Panthera leo* | 0.15 | 0.16 | 0.17 | 0.17 | 0.02 | 0.01 | | 0.03 | 0.13 | 0.17 | 0.23 | 0.26 | 0.21 | 0.19 | 0.24 | 0.26 | 0.27 | 0.27 | 0.27 | 0.42 |
| | 8 | *Leopardus geoffroyi* | 0.14 | 0.15 | 0.16 | 0.16 | 0.01 | 0.01 | 0.01 | | 0.11 | 0.15 | 0.21 | 0.24 | 0.18 | 0.17 | 0.23 | 0.26 | 0.26 | 0.25 | 0.26 | 0.41 |
| | 9 | *Canis lupus familiaris* | 0.15 | 0.17 | 0.18 | 0.17 | 0.10 | 0.10 | 0.11 | 0.10 | | 0.20 | 0.24 | 0.26 | 0.19 | 0.16 | 0.21 | 0.23 | 0.25 | 0.27 | 0.28 | 0.41 |
| | 10 | *Tupaia belangeri* | 0.14 | 0.15 | 0.16 | 0.16 | 0.16 | 0.16 | 0.17 | 0.16 | 0.23 | | 0.23 | 0.26 | 0.18 | 0.15 | 0.23 | 0.24 | 0.25 | 0.25 | 0.29 | 0.43 |
| | 11 | *Rhinolophus ferrumequinum* | 0.13 | 0.14 | 0.15 | 0.15 | 0.15 | 0.15 | 0.15 | 0.14 | 0.16 | 0.16 | | 0.20 | 0.16 | 0.22 | 0.28 | 0.30 | 0.30 | 0.33 | 0.34 | 0.43 |
| | 12 | *Pteropus alecto* | 0.14 | 0.16 | 0.17 | 0.16 | 0.18 | 0.17 | 0.17 | 0.17 | 0.18 | 0.19 | 0.12 | | 0.13 | 0.22 | 0.28 | 0.27 | 0.27 | 0.32 | 0.29 | 0.42 |
| | 13 | *Sus scrofa* | 0.17 | 0.18 | 0.19 | 0.19 | 0.18 | 0.18 | 0.18 | 0.17 | 0.21 | 0.20 | 0.18 | 0.21 | | 0.18 | 0.27 | 0.24 | 0.24 | 0.26 | 0.27 | 0.42 |
| | 14 | *Bos taurus* | 0.17 | 0.18 | 0.19 | 0.19 | 0.21 | 0.20 | 0.21 | 0.20 | 0.24 | 0.26 | 0.19 | 0.22 | 0.17 | | 0.19 | 0.26 | 0.28 | 0.25 | 0.27 | 0.39 |
| | 15 | *Tursiops truncatus* | 0.13 | 0.14 | 0.15 | 0.15 | 0.15 | 0.15 | 0.16 | 0.14 | 0.17 | 0.17 | 0.14 | 0.17 | 0.11 | 0.10 | | 0.28 | 0.30 | 0.31 | 0.30 | 0.46 |
| | 16 | *Marmota monax* | 0.15 | 0.16 | 0.18 | 0.18 | 0.21 | 0.21 | 0.22 | 0.21 | 0.22 | 0.20 | 0.21 | 0.20 | 0.22 | 0.25 | 0.19 | | 0.03 | 0.25 | 0.23 | 0.41 |
| | 17 | *Ictidomys tridecemlineatus* | 0.15 | 0.17 | 0.17 | 0.18 | 0.21 | 0.21 | 0.21 | 0.21 | 0.22 | 0.20 | 0.21 | 0.21 | 0.21 | 0.24 | 0.18 | 0.02 | | 0.26 | 0.24 | 0.41 |
| | 18 | *Mus musculus* | 0.22 | 0.23 | 0.24 | 0.24 | 0.25 | 0.25 | 0.26 | 0.25 | 0.28 | 0.27 | 0.23 | 0.26 | 0.28 | 0.32 | 0.25 | 0.24 | 0.24 | | 0.10 | 0.46 |
| | 19 | *Rattus norvegicus* | 0.22 | 0.22 | 0.23 | 0.24 | 0.24 | 0.24 | 0.24 | 0.24 | 0.28 | 0.27 | 0.22 | 0.24 | 0.27 | 0.30 | 0.25 | 0.22 | 0.23 | 0.09 | | 0.48 |
| | 20 | *Ornithorhynchus anatinus* | 0.36 | 0.36 | 0.37 | 0.37 | 0.37 | 0.37 | 0.37 | 0.37 | 0.41 | 0.42 | 0.37 | 0.40 | 0.38 | 0.42 | 0.36 | 0.41 | 0.41 | 0.46 | 0.46 | |

### Supplemental Figure 1

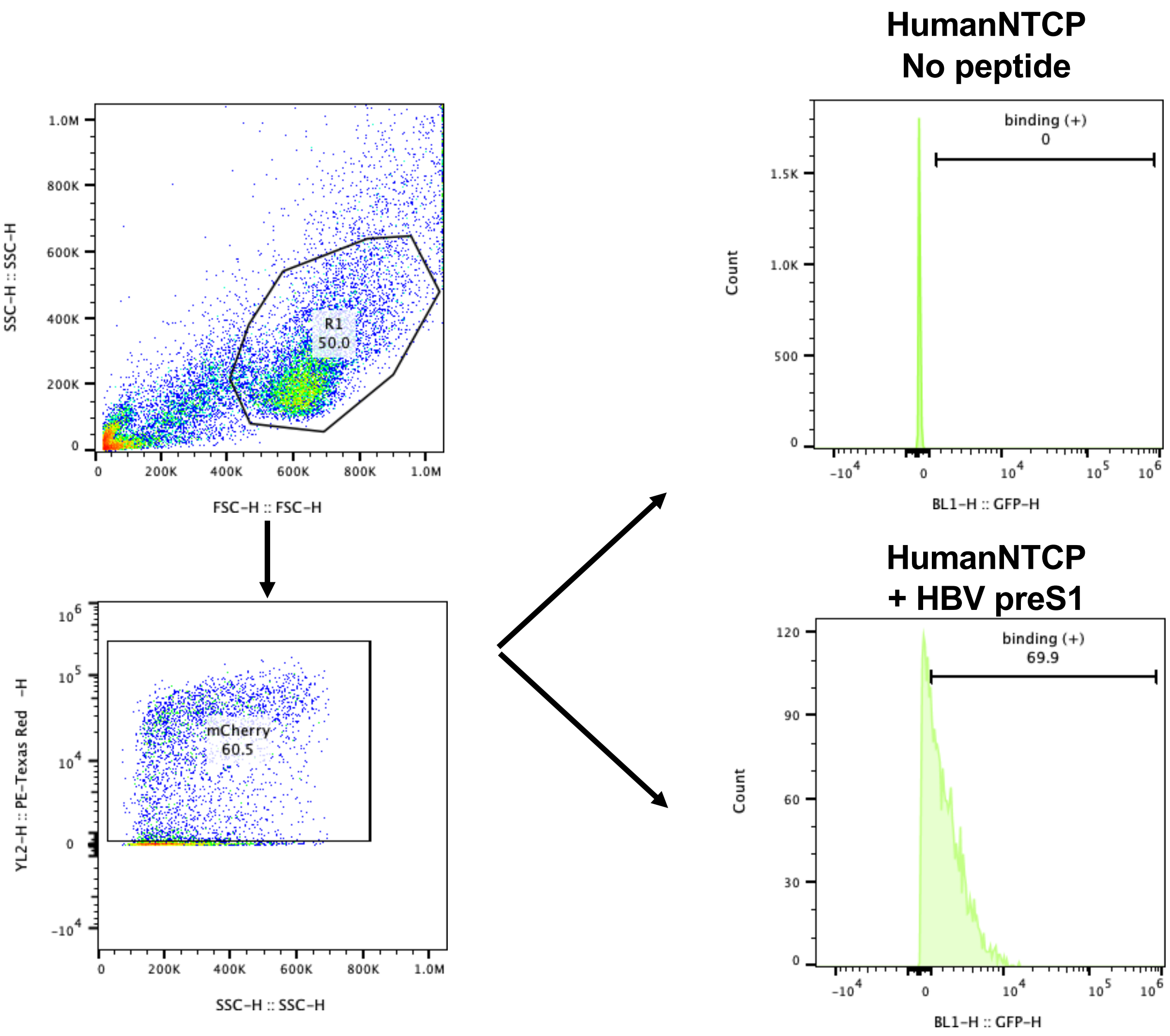

### Supplemental Figure 2

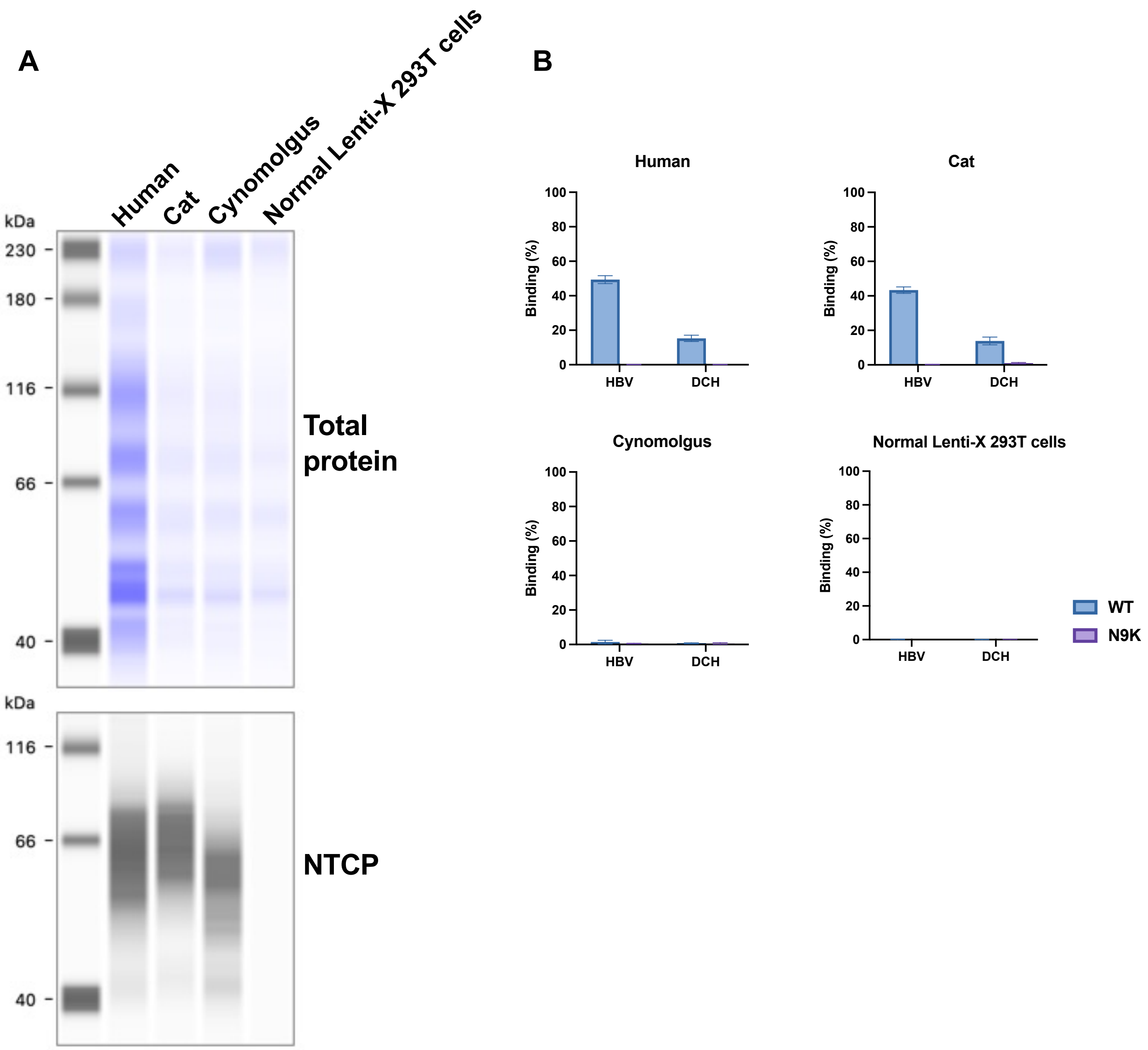

**Supplemental Figure 3**

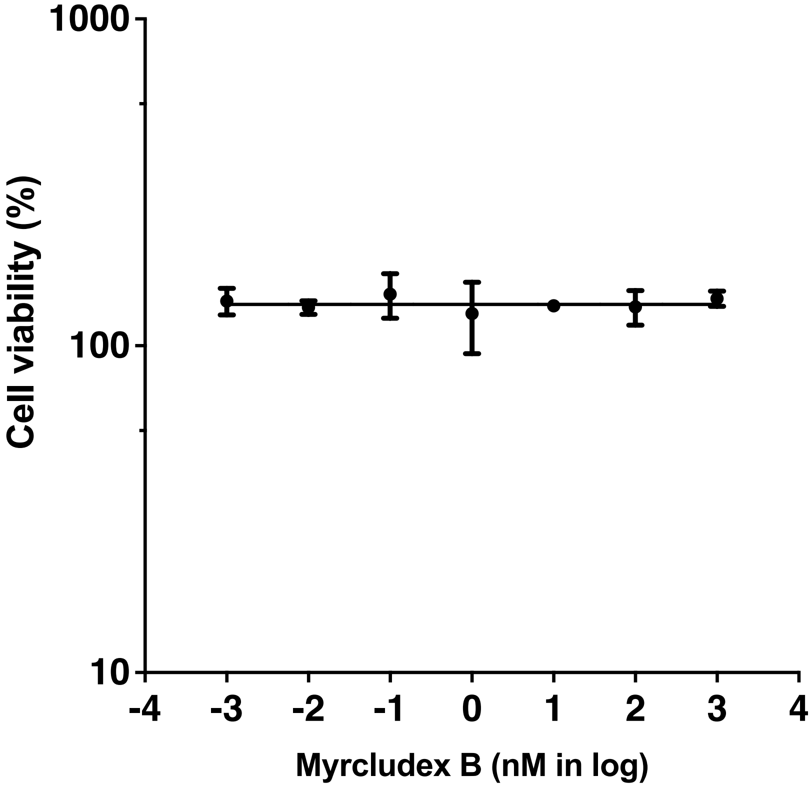
